# A new discrete-geometry approach for integrative docking of proteins using chemical crosslinks

**DOI:** 10.1101/2024.10.24.619977

**Authors:** Yichi Zhang, Muskaan Jindal, Shruthi Viswanath, Meera Sitharam

## Abstract

The structures of protein complexes allow us to understand and modulate the biological functions of the proteins. Integrative docking is a computational method to obtain the structures of a protein complex, given the atomic structures of the constituent proteins along with other experimental data on the complex, such as chemical crosslinks or SAXS profiles. Here, we develop a new discrete geometry-based method, wall-EASAL, for integrative rigid docking of protein pairs given the structures of the constituent proteins and chemical crosslinks. The method is an adaptation of EASAL (Efficient Atlasing and Search of Assembly Landscapes), a state-of-the-art discrete geometry method for efficient and exhaustive sampling of macromolecular configurations under pairwise inter-molecular distance constraints. We provide a mathematical proof that the method finds a structure satisfying the crosslink constraints under a natural condition satisfied by energy landscapes. We compare wall-EASAL with IMP (Integrative Modeling Platform), a commonly used integrative modeling method, on a benchmark, varying the numbers, types, and sources of input crosslinks, and sources of monomer structures. The wall-EASAL method performs better than IMP in terms of the average satisfaction of the configurations to the input crosslinks and the average similarity of the configurations to their corresponding native structures. The ensembles from IMP exhibit greater variability in these two measures. Further, wall-EASAL is more efficient than IMP. Although the current study uses crosslinks, the method is general and any source of distance constraints can be used for integrative docking with wall-EASAL. However, the current implementation only supports binary rigid protein docking, *i.e.*, assumes that the monomer structures are known and remain rigid. Additionally, the current implementation is deterministic, *i.e.*, it does not account for uncertainties in the crosslinking data beyond using distance bounds. Neither of these appears to be a theoretical or algorithmic limitation of the EASAL methodology. Structures from wall-EASAL can be incorporated in methods for modeling large macromolecular assemblies, for example by suggesting rigid bodies or restraints for use in these methods. This will facilitate the characterization of assemblies and cellular neighborhoods at increased efficiency, accuracy, and precision. The wall-EASAL method is available at https://bitbucket.org/geoplexity/easal-dev/src/Crosslink and the benchmark is available at https://github.com/isblab/Integrative_docking_benchmark.

## Introduction

Protein-protein interactions play a crucial role in biological processes, for example, in immune response, metabolism, growth, and development. Characterizing the structures of complexes formed by two or more proteins *via* a single experimental technique can often be challenging. Protein-protein docking methods aim to computationally determine the structure of the complex formed by two proteins, given their three-dimensional structures^1,2^. Some protein-protein docking methods employ rigid docking, where it is assumed that the proteins do not undergo significant conformational change upon binding. Rigid docking is computationally efficient since the search space is restricted to rigid translations and rotations of one protein with respect to the other. In integrative docking, additional experimental data, such as residual dipolar couplings from NMR spectroscopy and data from chemical crosslinking mass spectrometry (XLMS) can be used to guide the docking search^3–8^. Integrative docking methods aim to compute an ensemble of structures of the complex that are consistent with the experimental data. Here, we develop a new method for integrative rigid docking of protein pairs using inter-protein chemical crosslinks.

Chemical crosslinking involves treating the protein complex of interest with a chemical crosslinker^9,10^. A crosslinker consists of two reactive groups, separated by a spacer that defines the maximum crosslinker length. Common crosslinkers include DSSO (Disuccinimidyl sulfoxide), DMTMM (4-(4,6-Dimethoxy-1,3,5-triazin-2-yl)-4-methylmorpholiniumchloride), ADH (Adipic Dihydrazide), and EDC (1-ethyl-3-(3-dimethylaminopropylcarbodiimide hydrochloride) ^9,10^. The reactive groups in a crosslinker can bond covalently with accessible residues in a complex. Subsequent treatment with trypsin and analysis of the resulting mass spectrometry data provides a list of residue pairs in the complex that are crosslinked. Therefore, XLMS can provide upper bound distances between pairs of residues, which can inform the proximity of these residues in the structure of the complex.

Several methods exist for modeling protein complexes based on atomic structures and crosslinks^9–13^. Well-known integrative modeling methods such as Haddock, IMP, and Assembline can be used to dock proteins based on crosslinks^6,14–17^. These methods are broadly applicable, allowing for the use of diverse types of data, including data other than crosslinks and structures. Many of them also allow for determining the structure of larger complexes and assemblies. Some integrative modeling methods were developed specifically for modeling with crosslinking data, such as XLMOD^18^ and IMProv^19^. Recently, deep learning-based methods for structure prediction have been extended to use chemical crosslinks as additional inputs. Methods such as Alphalink, Alphalink2, and DistanceAF use the crosslinking data either in the input residue pair representation (Alphalink and Alphalink2) or in the loss function (DistanceAF)^20–22^.

Many of the aforementioned methods for modeling with crosslinks employ randomized sampling, *e.g.*, *via* Markov Chain Monte Carlo (MCMC) methods which is typically not stochastic, *i.e.*, the sampled regions of the landscape depend on the choice of initial random configurations of the sampling trajectories^17,23,24^. In particular, exhaustive sampling of complex landscapes is not guaranteed in these methods. Also, the combination of simplified coarse-grained bead representations with hard-sphere excluded volume restraints in some methods may make accurate modeling particularly challenging for protein structures with concavities, such as grooves and pits in the interfaces. These perceived limitations of current approaches motivated us to explore other sampling methods.

EASAL (Efficient Atlasing and Search of Assembly Landscapes) is a state-of-the-art discrete geometry-based methodology for roadmapping, sampling and analyzing the landscape of macromolecular configurations satisfying possible sets of pairwise inter-molecular distance constraints^25,26^. EASAL is both a resource-light, stand-alone method and also complements prevailing MC, MD and docking methods^27^, demonstrating superior performance, especially for discontinuous pair-potential energy landscapes. The EASAL methodology^26^ and curated open-source software^28^, https://bitbucket.org/geoplexity/easal-dev/src/master/ (see also http://www.cise.ufl.edu/~sitharam/EASALvideo.mpeg) can efficiently generate an exhaustive ensemble of structures lying within specific pair-potential wells, discretized as a staircase of nested distance-interval constraints. The methodology has been earlier used for effectively predicting virus assembly pathways^29^, sticky-sphere path integrals^26^ and free energy, configurational entropy, or volume computation^30,31^. Here, we leverage the unique features of the EASAL method for integrative docking with crosslinks. Given the structures of two proteins and an input set of crosslinks between them, the modified method, *wall-EASAL*, produces an ensemble of structures of the complex satisfying the maximum number of input crosslinks.

We compared the performance of wall-EASAL with IMP on the problem of integrative docking with crosslinks on thirty protein pairs, varying the number of input crosslinks, the crosslinker length, the source of crosslinks, and the source of monomer structures. Assessing the structure ensembles from these methods based on their average satisfaction of the input crosslinks and their average similarity to the corresponding native structures, we find that wall-EASAL performs better than IMP. It is also more efficient. Although the current study uses crosslinks, the method is general and any source of distance constraints can be used for integrative docking with wall-EASAL. The limitations are that the current implementation only supports binary protein docking, the monomer structures are assumed to be known and remain rigid, and the uncertainty in the crosslinking experiment is not considered. However, none of these appears to be a theoretical or algorithmic limitation of the EASAL methodology. Structures from wall-EASAL can complement methods for modeling large macromolecular assemblies by suggesting rigid bodies or restraints for use in integrative modeling methods^6,14,16,17,32–34^. This approach is expected to enhance the efficiency, accuracy, and precision at which large assemblies and cellular neighborhoods are characterized.

### Notes on terminology

We use (sampled) “structure of a complex” and “configuration” interchangeably. A “restraint” is a probabilistic term with biophysical origin referring to a constraint that may or may not be satisfied. A “constraint” is a geometric condition that is deterministic and Boolean. Either a constraint is satisfied, or it is not. We use “crosslink distance” to refer to the distance between crosslinked residues in a structure of the complex.

## Methods

### EASAL Background and Crosslink Satisfaction

EASAL is a geometric methodology designed to solve systems of distance interval constraints between point-sets, *i.e.,* to relatively position input point-sets with respect to each other so that they satisfy distance interval constraints between them. ^26^covers a general version of the algorithm in detail, yet a modified variant of the two-body version covered in^28^ suffices for this article.

#### The Computational Problem: Input, Assembly Constraint System and Desired Output

The input consists of the following:

⍰ A pair of finite point-sets (*A, B*) (sets of centers of *C_α_* atoms of a protein-pair, each protein assumed to be rigid), with all points in 3-dimensional Euclidean space;
⍰ A set *C* of distance interval constraints, where each constraint is between a pair of points in different point sets.

The unknown variable being constrained is a Euclidean isometry *T* applied to *B*, (*i.e.,* translation and rotation of *B* w.r.t. *A*). We call (*A,T*(*B*)) a *Cartesian configuration of* (*A, B*). The *assembly constraint system C* consists of 2 types of constraints: *C*_1_ being collision avoidance, and *C*_2_ being satisfaction of at least one crosslink constraint in a specified set cL of atom pairs. Formally,

⍰ 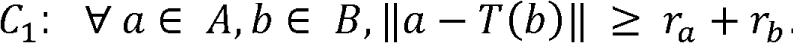. In other words, *T* is feasible only if (*A, T*(*B*)) is collision-free.
⍰ 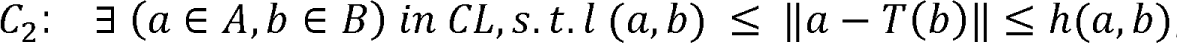, where 0 ≤ *l*(*a,b*) ≤ *h*(*a,b*). In other words, *T* is infeasible unless (*A, T*(*B*)) satisfies at least one of the given crosslinks. For this article, *l* and *h* are linear functions of *r_a_* + *r_b_* : *l*(*a, b*) = λ_*l*_(*r_a_* + *r_b_*) + δ_*l*_ and *h*(*a,b*) = λ_*h*_(*r_a_* + *r_b_*) + δ_*h*_.

The desired output is

⍰ a *feasible* Euclidean isometry *T* (resp. Cartesian configuration (*A, T*(*BB*))), defined as satisfying *C*_1_, *C*_2_, and simultaneously satisfying the maximum possible number *m* of crosslink constraints within *C*_2_, between pairs in *CL*, and
⍰ a representative sample set of feasible cartesian configurations (*A*, *T*(*B*)).

Generally speaking, solving such a system requires traversal of a 6-dimensional, topologically complex feasible Cartesian configuration space of Euclidean isometries T. Such traversal is impractical when the number of points in the point-sets and number of crosslinks is large. We simultaneously face the combined curses *of dimensionality and topological complexity*. EASAL mitigates these using the following strategies ^25,26,28,35–37^:

1. Generating an accurate atlas of the landscape, with minimal sampling, including a topological roadmap of configurational regions that satisfy different subsets of crosslink constraints in *CL* - called *active constraint regions (ACRs)* and formally defined in the next subsection – with a stratified neighborhood relationship between such regions. In other words, exploration is decoupled from sampling (Fig. 1).
2. Efficient sampling of ACRs, avoiding any form of gradient descent or retraction maps to enforce constraints, or repeated samples, and minimizing rejected samples. This is achieved by convexifying ACRs using a customized *Cayley parametrization,* introduced and developed over a decade ago ^26,28,36,37^, formally defined in the following subsections. This is a distance-based internal coordinate representation of configurations within ACRs, particularly suited to ACRs which are entirely defined using distance-based constraints (Fig. 1).

**Figure 1:**
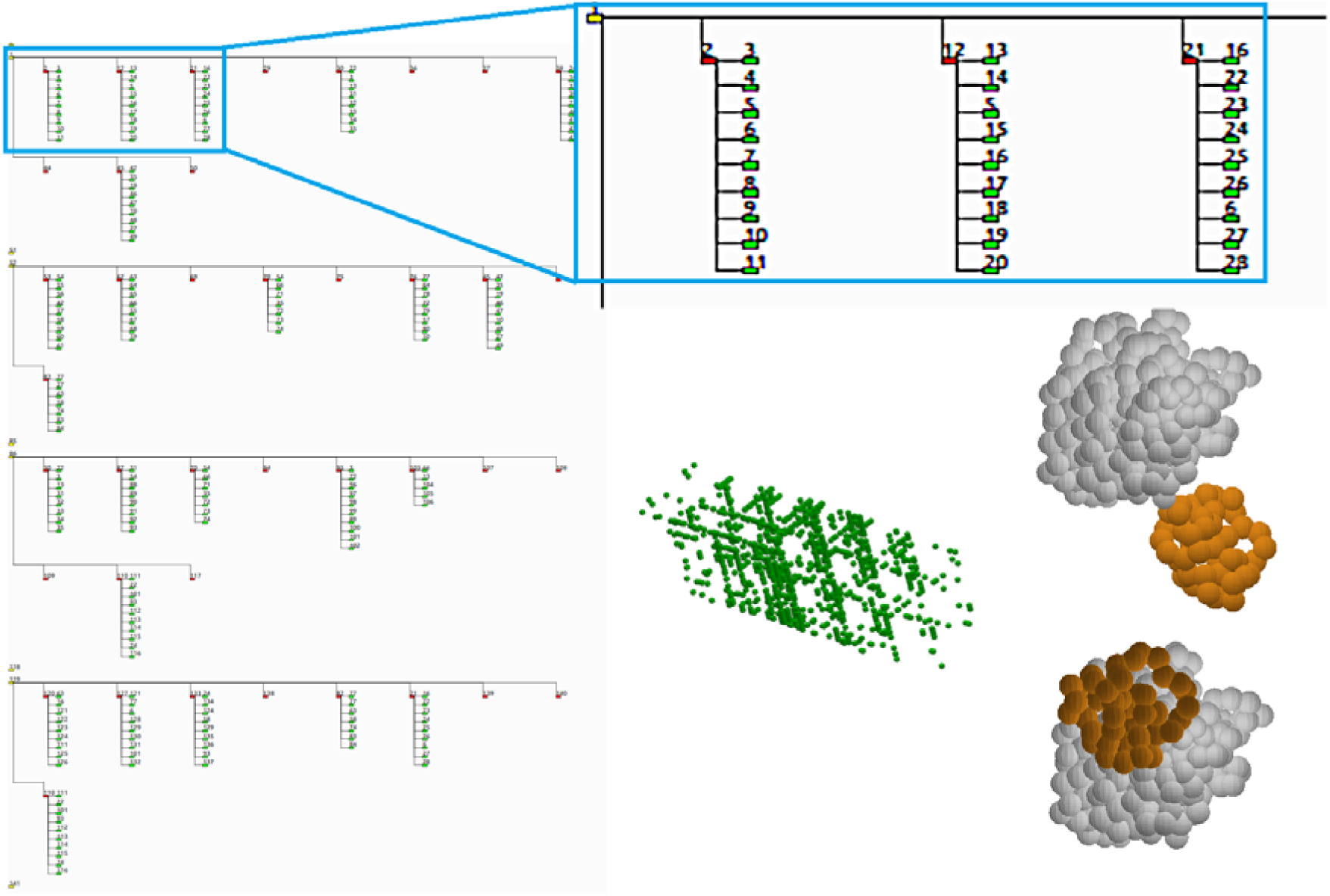
EASAL Background. Illustrative screenshots of EASAL from input case 1r0r/DSSO/3. Top: EASAL roadmap (directed acyclic graph represented as a tree by repeating nodes). Left: enlarged portion of the roadmap for detail; Bottom mid: view of the Cayley configuration space using distances between crosslinked residues as Cayley parameters, each green cube represents a feasible (collision-free and satisfying all crosslinks) configuration. Bottom right: selected configurations of the system satisfying all crosslinks and collision constraints, gray - Monomer A, orange - Monomer B.

In the next 3 subsections, we give a formal description of each of the above strategies and provide an outline of an overall algorithm directly following the existing EASAL methodology.

### Roadmap and Stratification of Active Constraint Regions

The problem of satisfying the constraints in the assembly constraint system *C* can be divided into subproblems associated with constraint systems related to *C* as follows. Notice that the existential quantifier in *C*_2_ can be replaced by a union over all simultaneously satisfied subsets *Q* ⊆ *CL* of crosslink constraints. Consider a variant of the assembly constraint system consisting of all collision-avoidance constraints in *C*_1_, but replacing *C*_2_ by the conjunction of crosslink constraints in a subset *Q*.

The *Active Constraint Region* (ACR) *R_Q_*^25,26,28^ is defined as the set of all configurations *T* satisfying all constraints in *C_1_* and *Q*. The set of ACRs is stratified by the size of *Q* and organized as a partial order (in the form of a directed acyclic graph (DAG)) called the *roadmap* (Fig. 1). Each node of the roadmap DAG represents an ACR, and directed edges (or parent-child relationships) reflect the superset-subset relationship of the ACRs, *R_Q_*, or alternatively the subset-superset relationship of the corresponding crosslink constraint sets *Q*. That is, a child node’s ACR is a subregion of its parent, and it always satisfies exactly one extra crosslink than its parent. Both the DAG and stratification structures of the roadmap are used for exploration by the algorithm outlined below.

### Cayley Parametrization

Assuming the crosslink distance interval constraints in the set *Q* have distinct lower and upper bounds *l* and *h*, the dimension of the ACR *R_Q* is potentially equal to the ambient dimension 6, and typically has a complex topology (Fig 1) arising from the combination of constraints in *C_1_* and *Q. Cayley parametrization*, formally defined below, was introduced and developed ^26,28–30,36,37^ over a decade ago. This parameterization is broadly applicable ^37–39^, and maps from *R_Q_* into a convex, and hence topologically simple base space of the same dimension, consisting of *Cayley configurations*.

Before formally defining the Cayley parametrization map, we motivate its use via three additional properties listed below, besides the key property of convexification. The combination of these properties ensures high sampling efficiency and accuracy, avoiding gradient-descent or retraction maps to enforce constraints, avoiding repeated sample configurations and minimizing discarded sample configurations.

⍰ The Cayley parametrization map from Cartesian ACR *R_Q_* to the convex base space is computationally easy (constant time), essentially just measuring distances within a cartesian configuration in *R_Q_* which have not been explicitly constrained.

⍰ The boundaries of the convex Cayley base space are easy to compute, and depend entirely on the crosslink constraints in *Q*. Additional collision avoidance constraints in *C_1_* carve out typically a small number of convex Cayley parameterizable regions from this convex Cayley base space.

⍰ The inverse (from Cayley to Cartesian) maps each *Cayley configuration* in the base space to at most 8 Cartesian configurations, each uniquely identifiable by chirality, and easily computable (constant time).

We now formally define *Cayley parametrization* ^25,26,28,36,37^.

Each ACR *R_Q_* has an underlying *active constraint graph* (ACG) *G_Q_* = (*V_Q_*,*E* ∪ *E_Q_*), whose vertex set *V_Q_* represents the set of the points involved in the crosslink pairs in *Q*, additionally ensuring inclusion of at least 3 vertices from each of the point sets *A* and *B*. There are two sets of edges:

⍰ (*E*): all pairs (*a_i_*, *a_j_*) and (*b_i_, b_j_*) for all *a_i_, a_j_* ∈ *A*, and (*b_i_, b_j_*)∈ *B*.

⍰ (*E_Q_*): pairs (*a, b*) ∈ *Q*.

Define a *nonedge* of a graph as a vertex pair not in its edge set. One way to represent or parametrize a Cartesian configuration (*A, T*(*B*)) in an ACR *R_Q_* is using the tuple of distances/lengths attained in the configuration by a subset *F* of nonedges of its ACG *G_Q_*. This tuple is the corresponding *Cayley configuration,* with each *Cayley coordinate or parameter* (value) of the tuple being a nonedge length (value). Formally, this defines an easily computable *Cayley parametrization map* π*_F_* that maps the ACR *R_Q_* into the Cayley base space π*_F_*(*R_Q_*). By choosing a minimal nonedge set F generically of size (6 - |*Q*|) whose addition makes the ACG π*G_Q* rigid ^35^, we ensure that a Cayley configuration corresponds to at most finitely many Cartesian configurations.

Next, given an ACR *R_Q_* we describe which ACG’s *G_Q_* have such a minimal, rigidifying set of nonedges *F*, so that the Cayley parametrization maps to a convex Cayley base (configuration) space π*_F_*(*R_Q_*). The formal theorem and proof can be found here ^26,28,36,37^, from which a simple algorithm follows for judiciously choosing Cayley parameters *F*. First, we define a relevant class of graphs.

A graph is a *complete 3-tree* if it can be obtained by an inductive construction that starts with a triangle graph and successively adds a new vertex adjacent to the vertices of a triangle in the graph constructed so far. A complete 3-tree has (3|*V*| - 6) edges and is minimally rigid in R^3^, therefore it generically has finitely many realizations.

We find a complete 3-tree *G*_Q_* that includes all the vertices in *V_Q_*, and 3 sets of edges: (1) *E_Q_* (2) a complete 3-tree with vertex set *V_Q_* ∩ *A*, and (3) a complete 3-tree with vertex set *V_Q_* ∩ *B*. Then the edges in *G*_Q_* that are not in any of the above 3 sets are chosen as the set *F* of Cayley parameters. It is shown in ^25,26^ that for almost all ACR’s, Cayley parameters satisfying these properties can be found, thereby ensuring a convex Cayley configuration (base) space.

### Algorithm Outline

Starting with an ACR *R_Q_* with *Q* being the empty set, recursively search for subregions or child ACRs *R_Q_*\* with *Q* ⊂ *Q*\*, satisfying successively more crosslinks. The recursive search is a standard depth-first search that generates the roadmap DAG.

Additional crosslink satisfaction is detected by sampling the convex Cayley (base) configuration space π*_F_*(*R_Q_*), where *F* is a set of nonedges of the ACG *G_Q_*, *i.e.,* convexifying Cayley parameters chosen in the manner described in the previous section. The sampling is in Cayley coordinates, *i.e.,* lengths of the nonedges in *F*.

As many as possible Cayley parameters in the set *F* are chosen from the specified crosslink pairs *CL* that are not always satisfied in *R_Q_*, *i.e.,* from *CL* \ *Q*. These add *a priori* boundaries to π_*F*_(*R_Q_*) being sampled.

For each sampled Cayley configuration, a constant time computation of π^−1^_*F*_ provides at most 8 pre-image Cartesian configurations. Further crosslinks that are not in *F* are detected *a posteriori* for each sampled configuration.

At a parent ACR *R_Q_*, while sampling π*_F_*(*R_Q_*), when a configuration satisfying one more crosslink is found, the child ACR *R_Q_*\* is created and sampling immediately begins on the child, until a leaf or sink ACR of the roadmap DAG is reached, in which all configurations are *feasible, i.e., satisfy the maximum number* m *of crosslinks, and are output by the algorithm*.

At the end of each round of depth-first search, we correct any missing ACR’s of the roadmap DAG caused by coarse sampling as follows. For any ACR in the roadmap, all for must exist as ancestors in the roadmap. Hence if an ancestor is missing, it is created, sampled, and a new depth-first search rooted at is originated.

The process naturally ends when all non-empty ACR’s have been sampled.

Overall, the methodology directly addresses the curse of dimension and complexity of landscapes while giving formal guarantees of elJciency, accuracy, robustness, and trade-olJs for the core algorithm.

Since EASAL’s roadmapping and sampling algorithm is integrally based on using inter-monomer residue-pair distances as both distance-interval constraints and Cayley parameters, EASAL is almost tailored for sampling configurations satisfying the crosslink (and collision) constraints. Maximal subset of crosslinks can be chosen that guarantee convexification of the corresponding Cayley configuration space. Further crosslinks outside this subset are checked *a posteriori* for each sampled configuration. However, as mentioned earlier, due to the relatively wide crosslink distance intervals, there is significant loss of efficiency in directly using EASAL for sampling configurations that satisfy all crosslinks.

### Wall-EASAL

To boost the efficiency that deteriorates when directly using EASAL to deal with a 6-dimensional region defined by the large distance intervals arising from the crosslink ranges, we develop a novel adaptation of EASAL, called *wall-EASAL* for reasons that will be clear from the discussion below. Wall-EASAL fully leverages the EASAL methodology’s ability to deal with *exact distance or small distance-interval* constraints, by mapping the constrained configuration spaces from their high ambient dimension to a convex Cayley space in their typically much lower intrinsic dimension. The advantage of the mapping is that it completely avoids the gradient descent used by prevailing methods to enforce distance constraints.

The key intuition behind wall-EASAL (mathematically proven in the Supporting Information, see Fig. 2) is that if the collision free configuration space is *path-connected*, and there is a feasible configuration satisfying all crosslink and collision constraints, then there must also exist relatively many feasible configurations in which some crosslink attains either its maximum or its minimum distance *exactly.* Such a configuration is called a *wall configuration*. This geometric principle called “concentration of measure” can be biophysically interpreted as follows: as the dimension of the region increases, the probability of configurations being close to the boundary increases rapidly, *i.e.,* the contribution to configurational entropy - by configurations close to the boundaries of configurational regions - increases rapidly with the region’s dimension.

**Figure 2:**
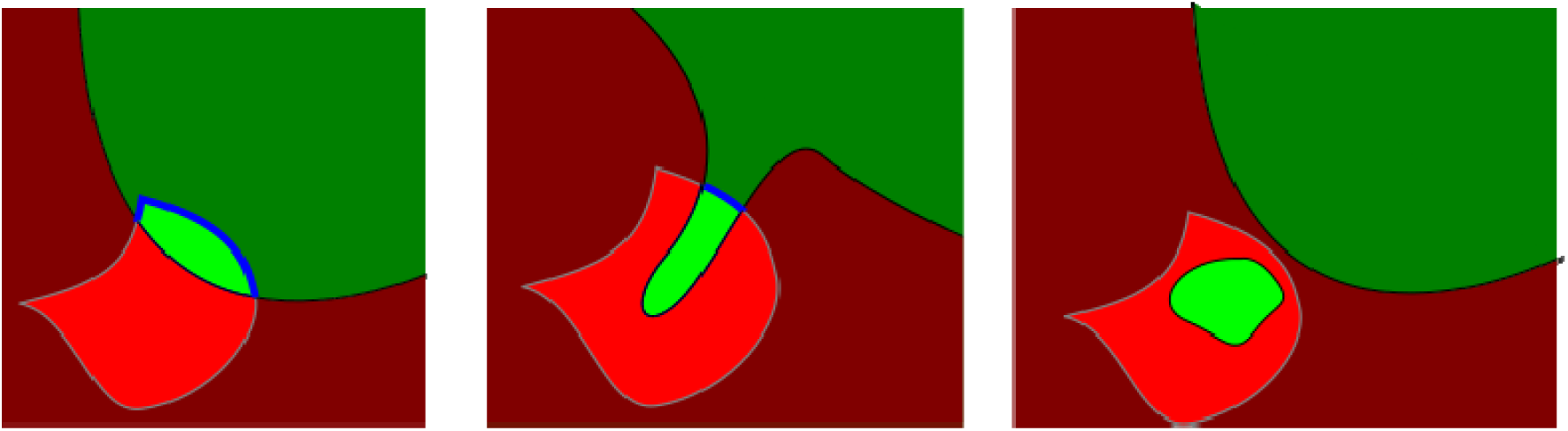
A schematic illustration of configuration spaces relevant to wall-EASAL. Shades of green: space of collision-free configurations, shades of red: space of colliding configurations. Lighter color: all crosslinks satisfied, darker color: not all crosslinks satisfied. The boundary between light and dark is a wall to which wall-EASAL sampling is restricted. Regular EASAL returns configurations in the light green region, wall-EASAL returns configurations in the blue curve only. Left: common/standard input cases; mid: input cases with a narrow bottleneck in collision-free configuration space; right: input cases disconnected with a pocket.

In the next 3 subsections, we employ this intuition to modify the direct “vanilla” application of the EASAL methodology described in the previous section.

#### Modified Computational Problem, Advantages

In addition to *C_1_* and *C_2_*, we introduce a new set of *wall* constraints, denoted *C*_3_. Specifically, we add to (or replace) every subset *Q* ⊆ *CL* of at most 6 crosslink constraints with a wall constraint set *Q_W_*:

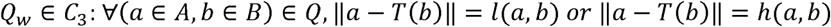

The key advantage of this modification is dimension reduction. Recall that the dimension of the ACR’s *R_Q_* in the previous section was potentially equal to the ambient dimension of 6 irrespective of the size of *Q*.

However, as formally described in the next 2 subsections, each wall constraint generically reduces the dimension of the ACR by 1: the dimension of *R_Qw_* is (6 - |*Q_W_*|), although the ambient Cartesian dimension remains 6.

Specifically, each 6-dimensional ACR in the previous approach is replaced by at most 2^|*Q_W_*^| (2^6^ = 64 but empirically on average no more than 4 nonempty) subregion ACRs *R_Qw_*, each being a (6 - |*Q_w_*|)-dimensional “wall” or boundary subset of the previous ACR *R_Q_*.

Therefore, the roadmap DAG is now additionally stratified by dimension, and although the ambient Cartesian dimension remains 6, the Cayley configuration space has equal intrinsic and ambient dimensions of (6 - |*Q_w_*|), while retaining the convexity property from the previous approach (when Cayley parameters are appropriately chosen). This mitigates the curse of dimensionality as well as topological complexity, making Cayley sampling of ACR’s computationally extremely efficient.

#### Modification of Roadmapping and Stratification

The introduction of wall constraints splits each ACR *R_Q_* in the previous approach into ACRs *R_Qw_*, each being a (6 - |*Q_W_*|)-dimensional “wall” or boundary subset of the previous ACR *R_Q_*. Therefore, each ACG *G_Q_* may correspond to multiple ACRs, no longer serving as a unique label. To resolve this issue and restore bijective labeling, we add a boolean label for each crosslink constraint in *Q*, *i.e.,* to each edge in *E_Q_*, indicating whether the minimum or the maximum value of that crosslink distance interval is enforced, so that each ACR *R_Q_w__* has a unique label specifying which wall of R_Q_ it corresponds to.

The parent-child relationship in the roadmap DAG is also altered. As in the previous section, in a parent-child pair, the ACGs differ by the inclusion of an extra edge; here we further require all their shared edges to have the same labeling.

As indicated by graph rigidity theory of the ACG *G_Q_*^35^, each extra wall constraint in *Q_w_* is typically independent, and therefore a child ACR is generically a one-dimension-lower-boundary of its parent ACR, with one extra crosslink distance fixed at one of the two extreme values, thereby relating the level of stratification of the roadmap DAG with the dimensionality of its ACRs. Hence, we can rely on an interior-to-boundary traversal and sampling procedure to populate the roadmap, as described in the next subsection.

#### Cayley Sampling and Boundary Searching

As mentioned, an ACR *R_Q_* in the wall-EASAL roadmap is a (6 - |*Q_w_*|)-dimensional object in 6-dimensional ambient space (Cartesian). Traditional methods of sampling would enforce |*Q_w_*| constraints in the ambient 6-dimensions, thus losing the advantage of the lower intrinsic dimension of *R_Q_w__* region and making the samspling computationally complex and challenging (see *e.g.*, left of Fig 3). Further illustrations can be found in ^26^ and cover page figure of ^27^.

**Figure 3:**
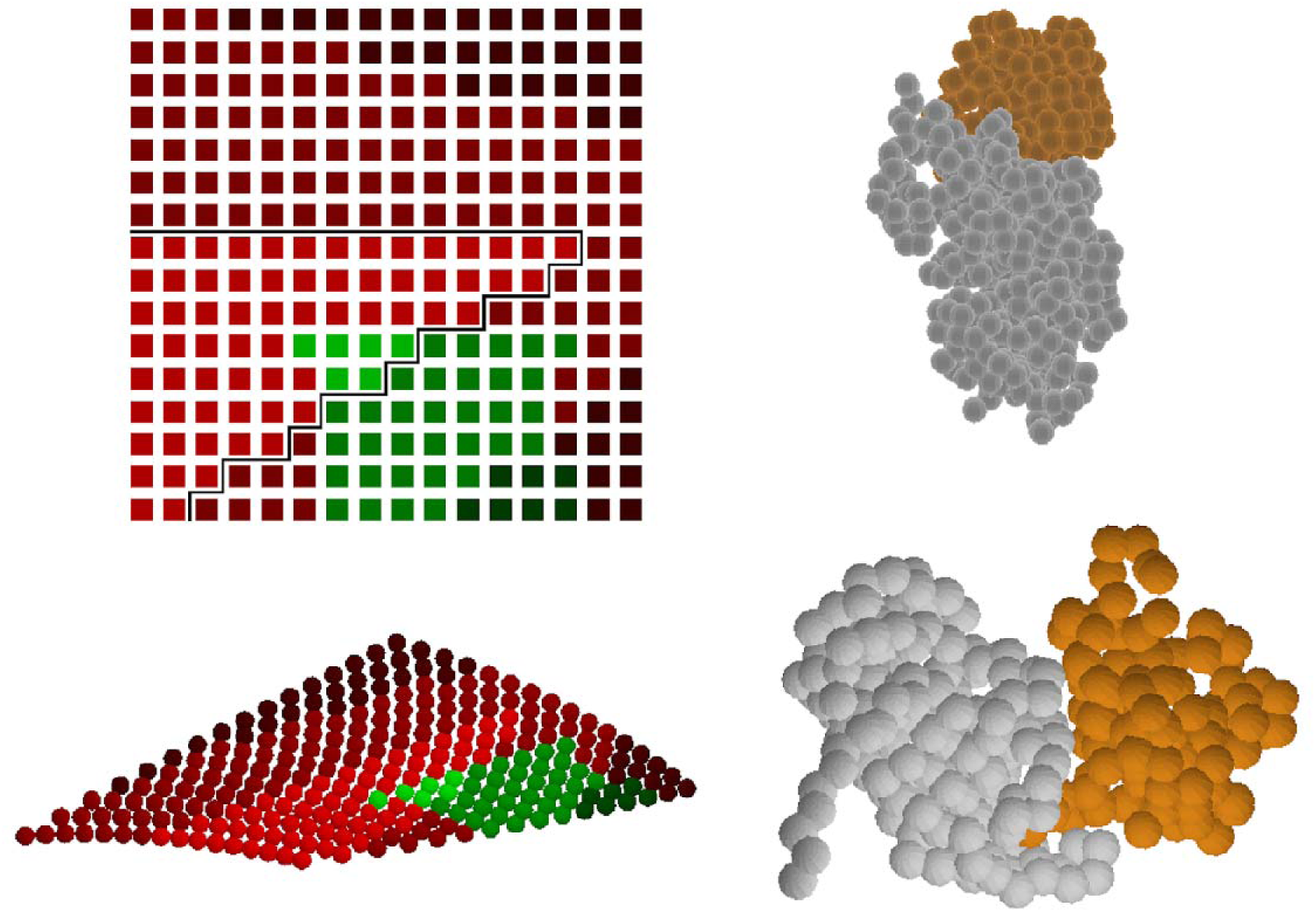
Pocket Artifact. Left: Cayley (above) and Cartesian (below) representations of a typical 2-dimensional slice of the configuration space for 2b42/DMTMM/10 in the neighborhood of a collision-free configuration (Top Right) satisfying all 10 crosslinks found by IMP. Wall-EASAL failed to find such a configuration due to pocket artifacts from *coarse* sampling. The slices were *finely* sampled for purposes of analysis/diagnosis. Each dot represents a sampled configuration, with Red = collision, Green = collision-free. Lighter shade of color means more crosslinks satisfied with the lightest being 10 (all crosslinks satisfied). Grey denotes the boundary of the region in which all 10 crosslinks are satisfied. Although the boundary between the lightest and slightly darker shade of green in fact consists of feasible wall configurations, since the entire feasible region is narrow, *coarse* sampling created a pocket artifact and caused wall-EASAL to miss this wall. However, for the input 2hle/DMTMM/9 with nearly the same number of crosslinks as 2b42, a variety of feasible configurations satisfying the maximum number of crosslinks (matching IMP) were found by wall-EASAL (Bottom Right).

As previously, we apply Cayley parameterization π*_F_* by choosing as Cayley parameters a minimal nonedge set *F* generically of size (6 - |*Q*|) whose addition makes the ACG *G_Q_* rigid^35^, ensuring that a Cayley configuration corresponds to at most finitely many Cartesian configurations and that the Cayley parametrization maps to a convex Cayley base (configuration) space π_*F*_(*R_Q_w__*) ^36^. The difference from previous “vanilla” EASAL is the following: while the dimension of π*_F_* (*R_Q_*) previously remained the same as the ambient dimension, *i.e.,* 6, now because the crosslink interval constraints in *Q* have been replaced by crosslink wall constraints, the dimension of π*_F_*(*R_Q_w__*) = |*F*| = 6 - |*Q*|. This is also the intrinsic dimension of a topologically complex *R_Q_w__* living in 6 ambient dimensions. Applying Cayley parametrization essentially “flattens” *R_Q_w__* to a simple, convex base Cayley configuration space whose ambient dimension is the same as its intrinsic dimension.

Furthermore, the discovery of a child ACR from a parent ACR is easier because of the boundary-interior relationship between these ACRs. When sampling π*_F_*(*R_Q_w__*) using Cayley coordinates, *i.e.*, lengths of nonedges in F, when a wall of an extra crosslink *c* ∈ (*CL* \ *Q*) is found, binary search can be used to find a configuration close enough to the wall or boundary where the crosslink c’s distance is at one of the end points of its interval. Then the child ACR *R_(QUC)_w__* is created in the roadmap, and sampling immediately begins on the child, until a leaf or sink ACR of the roadmap DAG is reached. Since the dimension drops by 1 from parent to child, for leaf ACR’s *R_Q_w__*,|*Q_w_*| ≤ 6. In all the leaf ACR’s, the remaining crosslink constraints in *CL* \ *Q_w_* are checked and *feasible configurations, i.e.,* those that satisfy the maximum number m of crosslinks, are output by the algorithm.

All other aspects of the previous algorithm outline remain unchanged.

Using this process, wall-EASAL returns all collision-free configurations *in the wall subset*, *i.e*., in which at least 1 crosslink has its distance at its wall, and all (or the maximum number of) crosslinks are satisfied.

If necessary, more walls can be introduced distributed in the interior of the crosslink distance-interval, *i.e*., between the two extremes of the interval, at the expense of reducing efficiency (see Discussion section). In this paper, however, wall-EASAL only uses the two extreme walls, which are shown to nevertheless provide a representative set of configurations.

The path-connectivity condition of the collision-free configuration is important for wall-EASAL to return the desired output of the original crosslink problem formulation (whose feasible configurations are not restricted to the wall subset). Otherwise, disconnectivity in the collision-free configuration space could result in the feasible configuration space – which is the intersection of collision-free and maximum crosslink-satisfying configuration spaces – lying in the interior of the latter as in Fig. 2 (right). In this case, no crosslink distance in such configurations lies at the extremes of the crosslink distance interval, and wall-EASAL will fail to find a feasible configuration, although one exists.

Next, we discuss two potential issues that could impact the performance of wall-EASAL. (1) How realistic is the path-connectivity assumption on the collision-free configuration space, which is crucial to guarantee that wall-EASAL finds a collision-free configuration that satisfies all crosslinks if one exists (or one that satisfies the maximum number of crosslinks)? (2) How representative are the wall configurations in the space of all (interior) feasible configurations?

### Walls and Pockets

Path-connectivity of the collision-free configuration space is guaranteed unless there is a collision-free configuration from which the two monomers are unable to untangle and break free while following a configurational path that avoids collisions. This scenario is avoided in most situations as long as (a) the monomers cannot be arranged into a *knot (*formally a *link* in topology terminology) configuration, and (b) there is no *pocket* in one monomer into which some part of the other monomer is jammed with no collision-free exit from the jammed configuration (Fig. 3).

Although true links and pockets are rare, such artifacts can arise from coarse sampling. *i.e*., although the collision-free configuration space may be path-connected, there could be a narrow bottleneck through which all paths must pass, effectively disconnecting coarsely sampled regions. Thus, the notion of path-connectivity that guarantees wall-EASAL’s accuracy is in fact a relative notion that depends on the coarseness of sampling (Fig. 3).

### Walls versus Interiors

Although the wall subset of configurations is non-empty provided the entire configuration space is either non-empty or path-connected, are walls representative of the entire feasible region including the interiors?

We answer the question affirmatively by providing quantitative results in the next section. The geometric intuition for the wall being representative of the whole is that the volume of a high-dimensional object (such as the feasible region in discussion here) mostly lies close to its boundary, or “most of a high-dimensional orange’s volume is at its peel”. This geometric principle called “concentration of measure” can be biophysically interpreted as follows: as the dimension of the region increases, the probability of configurations being close to the boundary increases rapidly, *i.e.,* the contribution to configurational entropy - by configurations close to the boundaries of configurational regions - increases rapidly with the region’s dimension.

Fig. 4 pictorially illustrates for the input case 1r0r/DSSO/3 how the wall subset is a good representation of the entire, much larger set of feasible configurations.

**Figure 4.**
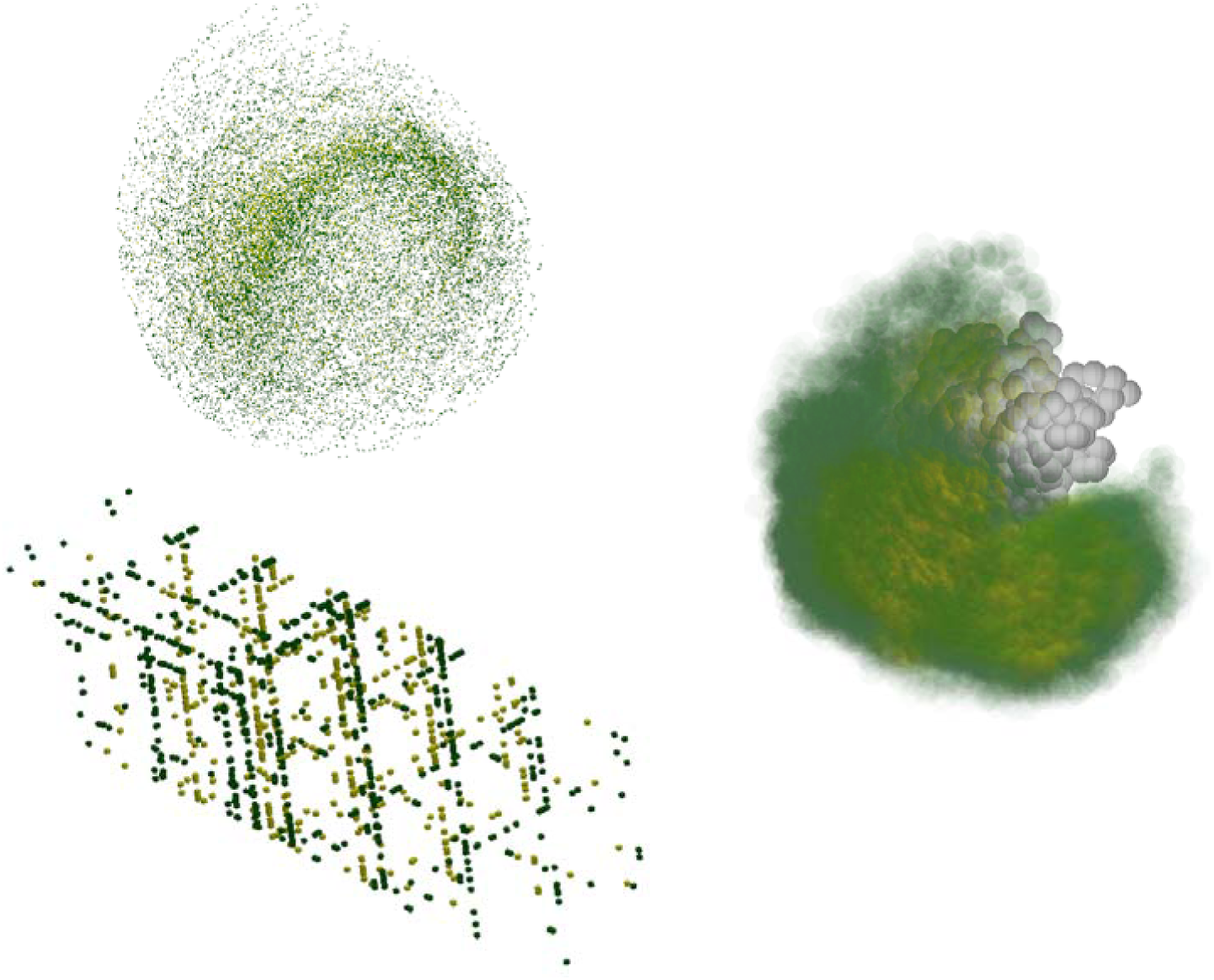
Representativeness of wall-EASAL sampling. Three different views of input case 1r0r/DSSO/3 showing all configurations on crosslink constraint walls (darker green) and not on walls (yellowish lighter green). Top Left: sampled configurations projected to 3 Cartesian dimensions (x, y, z), Bottom: projected to 3 crosslink distances used as Cayley parameters (the same as Fig. 1 Cayley configuration space). Right: sweep view of wall and interior feasible configurations of Monomer B (dark green - on wall, light green - off wall) with respect to Monomer A (gray) held fixed.

### Benchmark creation

#### Structures

We constructed a benchmark consisting of thirty integrative modeling cases of binary complexes (Table S1). There are twelve hetero-dimers, comprising seven complexes with experimentally solved structures from the Zlab 5.5 protein-protein docking benchmark^38^, and five AlphaFold-multimer predicted hetero-dimers from a crosslinking study^39^. Protein pairs with concave interfaces, containing grooves and pits were selected for docking, as these are perceived to be more difficult to dock than those with flat interfaces (Fig. S1). The monomer structures from the bound structure of the complex were used as input to the docking methods; the bound and unbound structures of the monomers are very similar for these cases.

#### Crosslinks

We generated two kinds of crosslinks: a longer crosslinker between lysines (DSSO) and a shorter crosslinker between aspartic acid and glutamic acid residues (DMTMM). DSSO and DMTMM crosslinks were simulated using Jwalk^40^. The maximum distances between crosslinked residues in Jwalk were set to 32 Å (20 Å) for DSSO (DMTMM)^41^. A false positive rate of 20% was used (default in Jwalk). Random subsets of inter-protein crosslinks from Jwalk were used for the benchmark cases.

In all, the benchmark consisted of twenty-five input cases with simulated crosslinks on experimentally determined structures of complexes from the Zlab 5.5 benchmark; the number and type of input crosslinks were varied across these cases. Additionally, there were five input cases with crosslinks from experimental studies alongside AlphaFold-multimer predictions of the complexes (Table S1).

### Running IMP on the benchmark

We used the Integrative Modeling Platform’s Python Modeling Interface (PMI) (IMP 2.17.0; https://integrativemodeling.org) for integrative docking. The modeling protocol was adapted from previous studies^6,17,23,42,43^.

The monomers were represented as independent rigid bodies based on their structures and coarse-grained at one residue per bead centered at the C_α_ atom. Bayesian crosslinking restraints were used, along with excluded volume and sequence connectivity restraints ^17,23,44^. The Gibbs sampling Replica Exchange Markov Chain Monte Carlo (MCMC) algorithm was used for structural sampling. We started with initial random configurations for each protein in each pair. A configuration, *i.e.,* structure of the complex, was saved after every ten Gibbs sampling steps, each of which consisted of a cycle of Monte Carlo moves comprising random translations and rotations of the monomer rigid bodies. We performed twenty independent runs with four replicas and ten thousand MCMC steps per run, resulting in eight million configurations for each benchmark input. Following this, we applied the analysis and validation protocols used in previous studies^17,23,45^.

### Running wall-EASAL on the benchmark

In wall-EASAL, each monomer was similarly represented as a rigid body, coarse-grained at one residue per bead, where each bead was centered at the C_α_ atom and had a radius of 2.5 Å. Each crosslink was encoded by two distance constraints (‘walls’), one corresponding to the upper bound and another corresponding to the lower bound on the distance between the centers of C_α_ atoms of the crosslinked residues. The lower bound distance for both DMTMM and DSSO crosslinks was set to 10 Å based on the lengths of the crosslinked lysine and glutamic/aspartic acid side chains and the rigid parts of the crosslinker. The upper bound distance was set to 32 Å (20 Å) for DSSO (DMTMM) crosslinks based on the linker and side chain lengths and the backbone flexibility of the monomers^41,44,46,47^. For each crosslink, two active constraints were defined in wall-EASAL, corresponding to the lower and upper bounds of the crosslink. An active constraint in wall-EASAL is of the form λ * (*r_i_* + *r_j_*) + δ, where λ and δ are constants and *r_i_* and *r_j_* correspond to the radii of the crosslinked beads^26,28,35^. Based on the above crosslink bounds, the values of λ and δ were set to 2 and 0 respectively for the lower bound constraint, and 0 and 32 (or 20) for DSSO (DMTMM) for the upper bound constraint. For all crosslink constraints, *r_i_* = *r_j_* = 2.5 Å was used. The threshold for the number of crosslinks to be satisfied by a configuration (‘*crossLinkSatisfyThres*’) was set to *n*-2 where n is the number of crosslinks. The collision constraint, similar to the excluded volume restraint in IMP, was used to avoid overlapping of beads. The parameters for the collision constraints, λ and δ, were set to 1 and 0, respectively. Lastly, we used a step size of 5, which defines the resolution of sampling.

### Analyzing wall-EASAL and IMP configurations

The ensemble of configurations from wall-EASAL and IMP were compared based on their crosslink satisfaction and similarity to the corresponding native structures on the benchmark. The crosslink satisfaction for an ensemble was determined based on two measures: the maximum percentage of crosslinks satisfied by any configuration in an ensemble and the average distance between crosslinked residues across all crosslinks and configurations in the ensemble. A crosslink was satisfied by a configuration if the distance between the bead centers, corresponding to the crosslinked residues, was within an upper bound of 32 Å (20 Å) for DSSO (DMTMM) crosslinks.

The native structure for each complex was the experimentally determined PDB structure of the complex (first twenty-five cases in Table S1) or the AF-multimer prediction (last five cases in Table S1). The similarity of the configurations to the corresponding native structure was determined by two measures: the distance between crosslinked residues difference and the ligand RMSD (root-mean-square deviation). The former is the difference between the distance between crosslinked residues in a sampled configuration to the corresponding distance in the native structure across all crosslinks and across all configurations in an ensemble. The ligand RMSD between a configuration and the native structure was calculated as the C_α_ RMSD of the second protein (ligand) after superposing the first protein (receptor) in the complex ^1,2^.

## Results

Here, we compare and contrast the performance of two methods for integrative docking of protein pairs using chemical crosslinks. We first demonstrate that the new method, wall-EASAL, which samples the configurations at the wall, is representative, by comparing it with the vanilla EASAL which performs exhaustive sampling in the entire search space under distance constraints which we call *interior-EASAL,* in order to clearly differentiate the methods. Next, we compare and contrast the performance of wall-EASAL with that of IMP on a benchmark set of input cases. This comparison is based on the crosslink satisfaction of the respective configurations, the similarity of the configurations to the native structures, and the efficiency of the methods. Finally, we visualize the structures of the complex produced for a few input cases.

### Coverage of the Interior using Wall-EASAL

To show that the ensemble generated by wall-EASAL sampling is indeed representative of the entire region of configurations satisfying constraints, we ran a coverage test similar to ^31,48^ by sampling both the entire region (using interior-EASAL) and the walls only (using wall-EASAL) and comparing these results to see if the smaller sample set “covers” the larger one. The coverage experiment is designed as follows.

For each input case, both sampling methods were executed yielding 2 sets of feasible configurations. Then the entire configuration space was partitioned into a 6-dimensional hypercube grid, and points generated by interior-EASAL were mapped into the hypercubes. We iteratively coarsened the grid (making each grid cube larger) until at least 90% of the occupied cubes had at least r sampled points in them. Here the coefficient is defined as r = Cinterior/wall)^1/6^ to avoid bias caused by difference in number of points between sampling methods. After grid size was determined, points sampled with wall-EASAL were mapped into such a grid, and all cubes with at least r interior samples were checked to determine if they have at least 1 wall sample in it. The ratio between numbers of those cubes was then tabulated.

Since wall volume is expected to be a lower percentage of the larger interior volume when there are fewer crosslinks, we ran four representative cases with few crosslinks (2, 3, 4, and 5, respectively) to demonstrate the representativeness. In all these cases, wall-EASAL provides an ensemble as good as EASAL in terms of coverage. Specifically, the coverage rate of wall-EASAL is constantly over 80%, corroborating our conclusion that sampling only the walls of the feasible region provides a good coverage of the entire feasible region including the interior.

**Table 1:** Coverage result of wall-EASAL. Number of sampled configurations in each method and grid cubes they are mapped to are tabulated here.

| Crossli<br>nker | PDB | Number of<br>crosslinks | Coverage<br>ratio | Interior sample<br>count | Wall sample<br>count | Interior<br>volume | Wall<br>volume |
| --- | --- | --- | --- | --- | --- | --- | --- |
| DSSO | 1clv | 2 | 0.84 | 13604 | 2941 | 192 | 163 |
|  | 1r0r | 3 | 0.97 | 36105 | 5305 | 81 | 79 |
|  | 2ayo | 4 | 0.97 | 108092 | 10444 | 172 | 168 |
|  | 2hle | 5 | 1 | 128 | 128 | 51 | 51 |

### Percentage of crosslinks satisfied

We first examined the number of crosslinks satisfied by the IMP and wall-EASAL ensembles in the thirty input cases in the benchmark. An integrative structure satisfies an input crosslink if the corresponding Cα-Cα distance between the crosslinked residues is less than the upper bound; the upper bounds depend on the linker lengths and were set to 32 Å (20 Å) for DSSO (DMTMM) linkers^7,17,23^.

The wall-EASAL and IMP configurations satisfy the crosslinks similarly well in terms of the highest percentage of crosslinks satisfied by a single configuration in the ensemble (Fig. 5A, Fig. S2). However, the distributions of crosslink percentages in the ensembles suggest that the wall-EASAL configurations satisfy a greater percentage of crosslinks on average (Fig. 5B-5E, Fig. S2). The IMP ensemble is more diverse in terms of crosslinks satisfaction.

**Figure 5.**
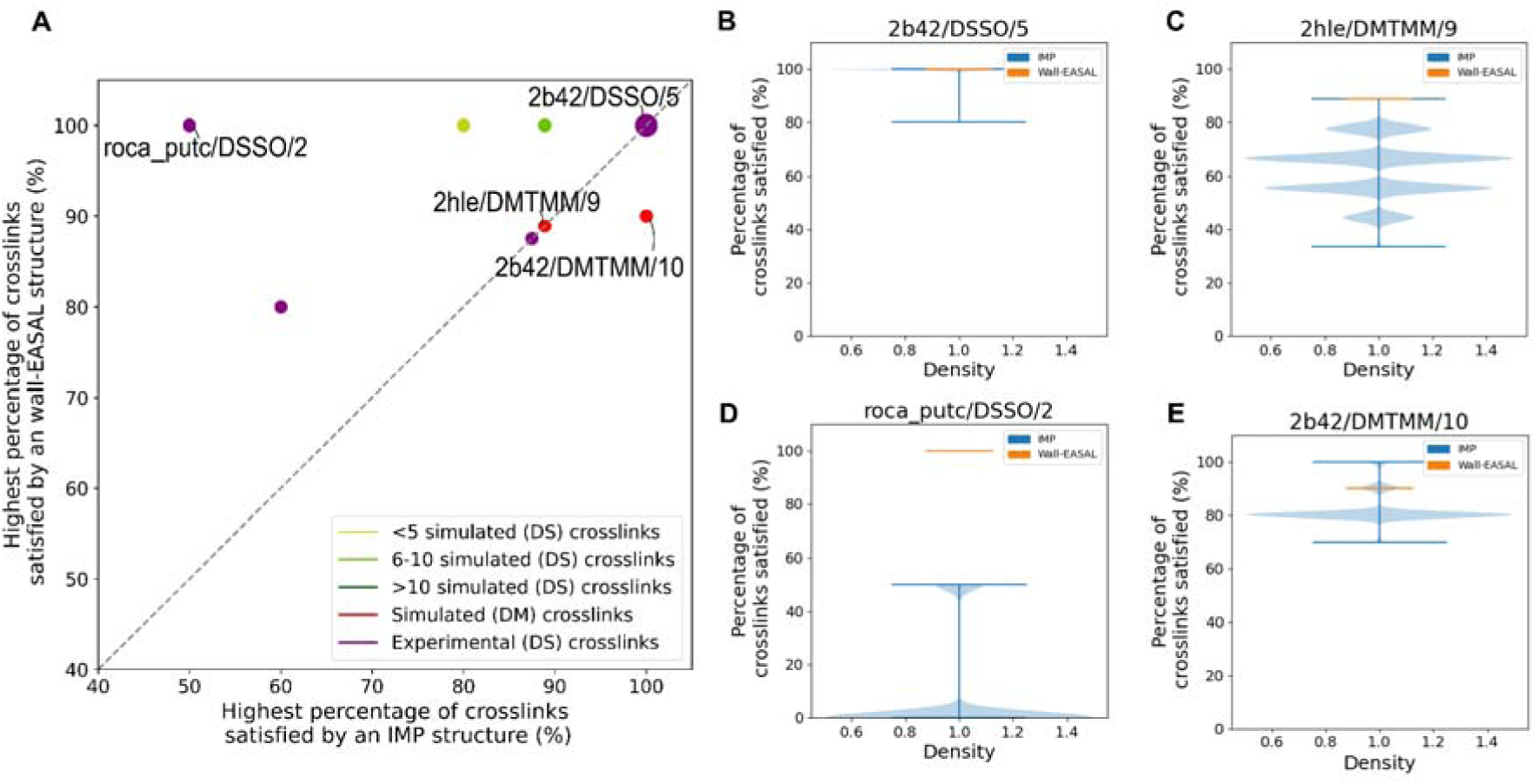
Percentage of crosslinks satisfied in wall-EASAL and IMP structures. For 30 benchmark inputs, the wall-EASAL (orange) and IMP (blue) ensembles are compared. **(A)** The highest percentage of crosslinks satisfied in any configuration in both the ensembles, where the larger point (top right) depicts the majority of the cases in which at least one configuration satisfies all the crosslinks. **(B-E)** Distribution of the percentage of crosslinks satisfied per configuration in the two ensembles for four cases. DS and DM refer to DSSO and DMTMM crosslinks, respectively.

We provide a few examples. In most input cases, both the wall-EASAL and IMP configurations satisfy the crosslinks equally well. For example, in 2b42/DSSO/5 all the wall-EASAL and IMP configurations satisfy all the crosslinks (Fig. 5B, Fig. S2), and in 2hle/DMTMM/9, the highest percentage of crosslinks satisfied by a configuration is 88% in both ensembles (Fig. 5C, Fig. S2). However, in a few input cases, wall-EASAL ensembles satisfy more crosslinks. For example, in roca_putc/DSSO/2, the highest percentage of crosslinks satisfied by an IMP configuration is only 50%, whereas the wall-EASAL configurations satisfy all the crosslinks (Fig. 5D, Fig. S2).

A sole exception is 2b42/DMTMM/10, in which an IMP configuration satisfies more crosslinks (100%) than a wall-EASAL configuration (90%) (Fig. 5E, Fig. S2). In this case, wall-EASAL performed slightly inferior to IMP primarily because of Wall-EASAL experiments’ coarse sampling. As pointed out earlier, the resulting pocket or disconnectivity artifacts in the collision-free configuration space cause wall-EASAL to miss wall regions whose projection on the chosen Cayley parameters (*i.e.*, inter-monomer residue-pair distances) is narrower than sampling step size.

For IMP, we observe that increasing the number of crosslinks improves the performance of integrative docking. The more input crosslinks, the higher the percentage of crosslinks satisfied per configuration, as shown by the shift in the blue distributions to higher crosslink percentages (Fig. S2A-S2C). This could indicate that, in randomized sampling guided by restraints, increasing the quantity of input data facilitates the sampling of more good-scoring configurations (configurations consistent with the input data). In contrast, wall-EASAL’s performance is largely independent of the number of crosslinks.

### Average crosslink distance

Next, we computed the distances between the crosslinked residues (crosslink distances) in the two ensembles. The wall-EASAL configurations satisfy crosslinks better in terms of the crosslink distances (Fig. 6, Fig. S3). In 29 (19) cases, the average crosslink distances were within the upper bounds in the wall-EASAL (IMP) ensembles (Fig. 6A). All wall-EASAL crosslink distances were usually within the upper bounds; in contrast, a fraction of the IMP configurations violated the crosslinks in all cases (Fig. S3). The range of crosslink distances is smaller for wall-EASAL and larger for IMP (height of violin plots, Fig. S3).

**Figure 6.**
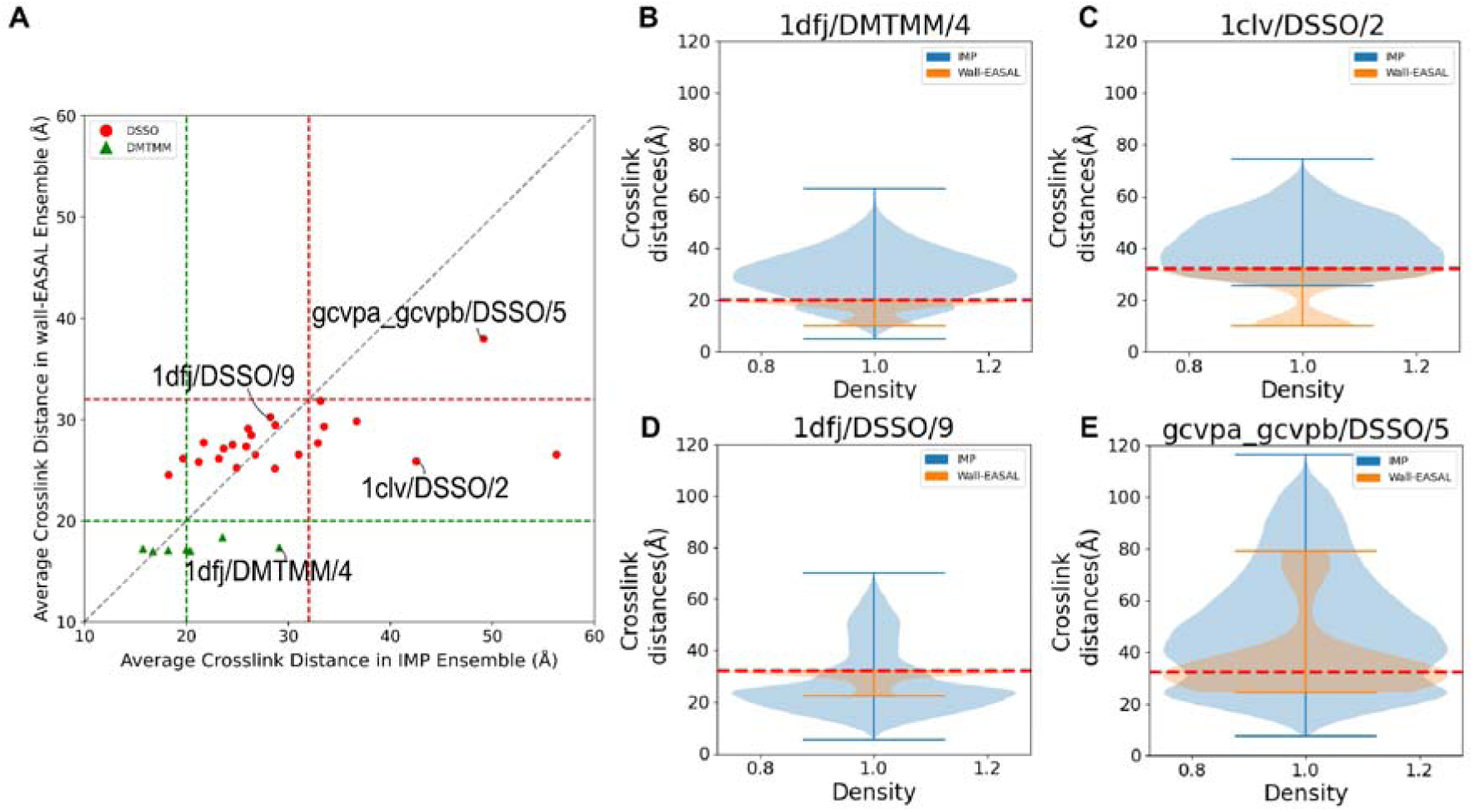
Distances between crosslinked residues in wall-EASAL and IMP structures. **(A)** For the 23 cases with DSSO crosslinker (red circles) and 7 cases with DMTMM crosslinker (green triangles), the average distances between the crosslinked residues were computed across all crosslinks and all configurations in the wall-EASAL and IMP ensembles. The crosslink upper bound (red and green dashed line) was set to 32 Å (20 Å) for DSSO (DMTMM) crosslinks. **(B-E)** The distribution of the distance between crosslinked residues in the two ensembles is shown for four cases.

In many input cases, the average crosslink distance was within the upper bound for both ensembles, *e.g*., 1dfj/DSSO/9 (Fig. 6B). However, for some cases, such as 1clv/DSSO/2 and 1dfj/DMTMM/4, the average crosslink distance exceeded the upper bound in the IMP, but not in the wall-EASAL ensemble (Fig. 6C-6D, Fig. S3). Finally, there were a small number of input cases, such as gcvpa_gcvpb/DSSO/5, where the average crosslink distance was much higher than the upper bound in both ensembles (Fig. 6E).

In most wall-EASAL configurations, the crosslink distances were close to the upper bound, especially as the number of crosslinks increased (Fig. S3A-S3C). This is unsurprising as wall-EASAL explicitly samples and finds only configurations where at least one crosslink distance takes an extreme value at one of the endpoints of its allowed interval.

Finally, for IMP, the crosslink distances reduce with an increase in the number of crosslinks, consistent with the decrease in docking difficulty with an increase in the number of crosslinks as shown earlier (Fig. S3A-S3C).

### Comparison of crosslink distances: sampled configuration vs native structure

Further, we compared the crosslink distances in the wall-EASAL and IMP configurations with the corresponding distances in the native structure. The crosslink distances in the wall-EASAL configurations were closer to those in the native structure, implying that the wall-EASAL ensemble contained more near-native configurations (Fig. 7A, Fig. S4). The differences between crosslink distances were more variable for the IMP ensemble (Fig. 7B-7E, Fig. S4).

**Figure 7.**
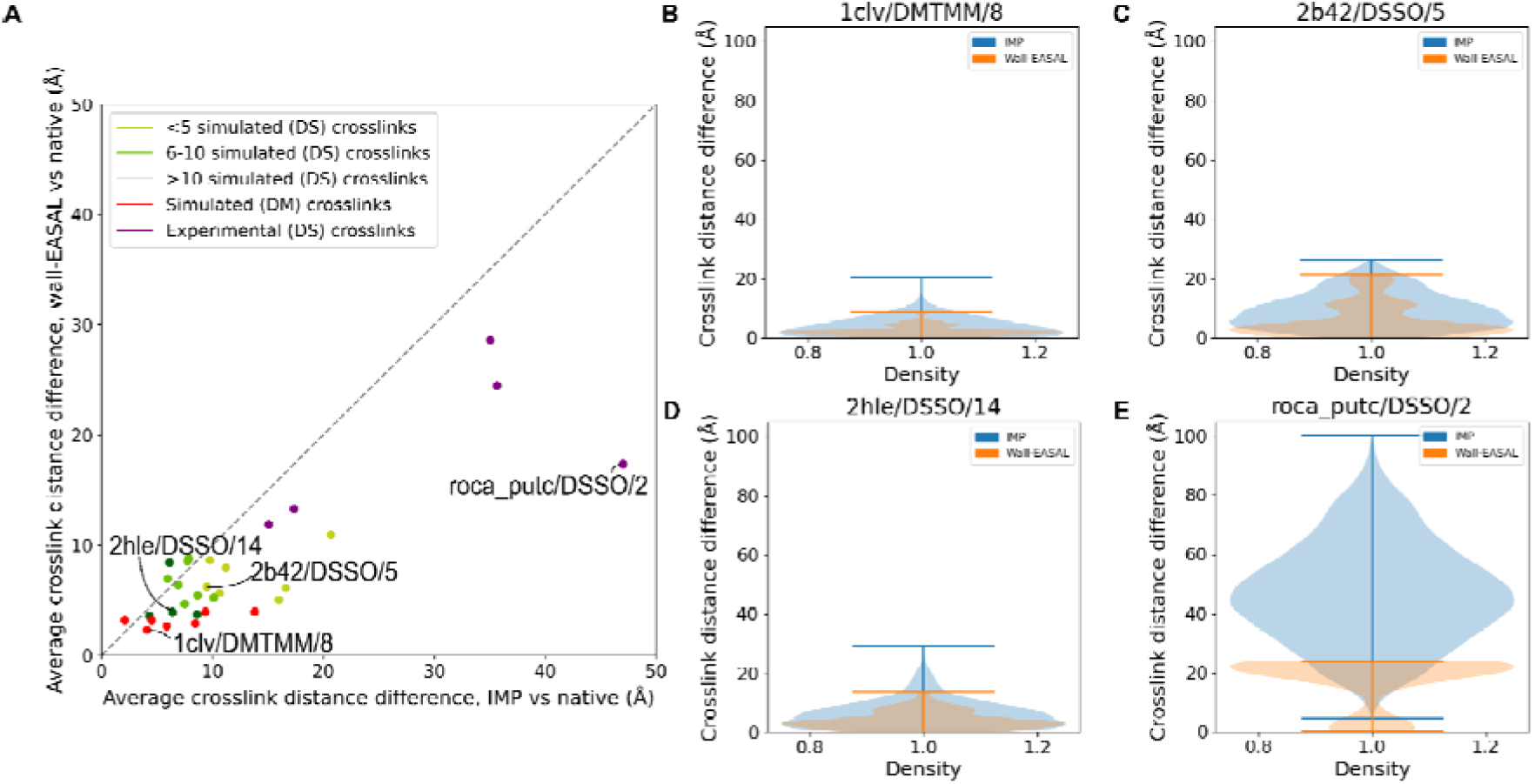
Comparison between crosslink distances in the sampled configurations and the native structure. **(A)** The average difference between the crosslink distance in a sampled configuration and the crosslink distance in the native structure was compared for the IMP and wall-EASAL ensembles. **(B-E)** The difference in crosslink distance between the native structure and each configuration in the two ensembles for four cases. DS and DM refer to DSSO and DMTMM crosslinks, respectively.

For most input cases with simulated DMTMM crosslinks, the crosslink distances from both the wall-EASAL and IMP configurations were close to those in the native structure, *e.g.*, 1clv/DMTMM/8 (Fig. 7B). For input cases with simulated DSSO crosslinks, the distances from the wall-EASAL configurations were closer to those in the native structure, e.g., 2b42/DSSO/5 and 2hle/DSSO/14 (Fig. 7C-7D). The differences in crosslink distances in both sets of configurations were larger for the input cases with crosslinks from experiments and AF2-predicted monomer structures, *e.g.*, roca_putc/DSSO/2, implying that these cases were more difficult for both methods (Fig. 7E). An increase in the number of crosslinks was associated with smaller distance differences in the IMP configurations, consistent with the earlier trends (Fig. S4A-S4C).

### Accuracy of the IMP and wall-EASAL configurations

We also computed the ligand RMSDs (Root-Mean-Square Deviation) of the configurations in the IMP and wall-EASAL ensembles with respect to the corresponding native structures^1,2^. Both methods performed similarly in recovering a single near-native configuration in the ensemble; however, the configurations in the wall-EASAL ensemble were closer to the native structure, on average (Fig. 8, Fig. S5). For 13 (12) input cases for IMP (wall-EASAL), the ligand RMSD of the best configuration was within 10 Å of the native structure (acceptable by CAPRI standards) (Fig. 8A). The average ligand RMSD to the native structure was 30 Å (38 Å) for wall-EASAL (IMP) (Fig. 8B). Configurations from both ensembles had higher RMSDs for the input cases with crosslinks from experiments and AF2-predicted monomer structures, *e.g.*, phes_phet/DSSO/5, consistent with the higher docking difficulty for these cases noted earlier (Fig. S5E).

**Figure 8.**
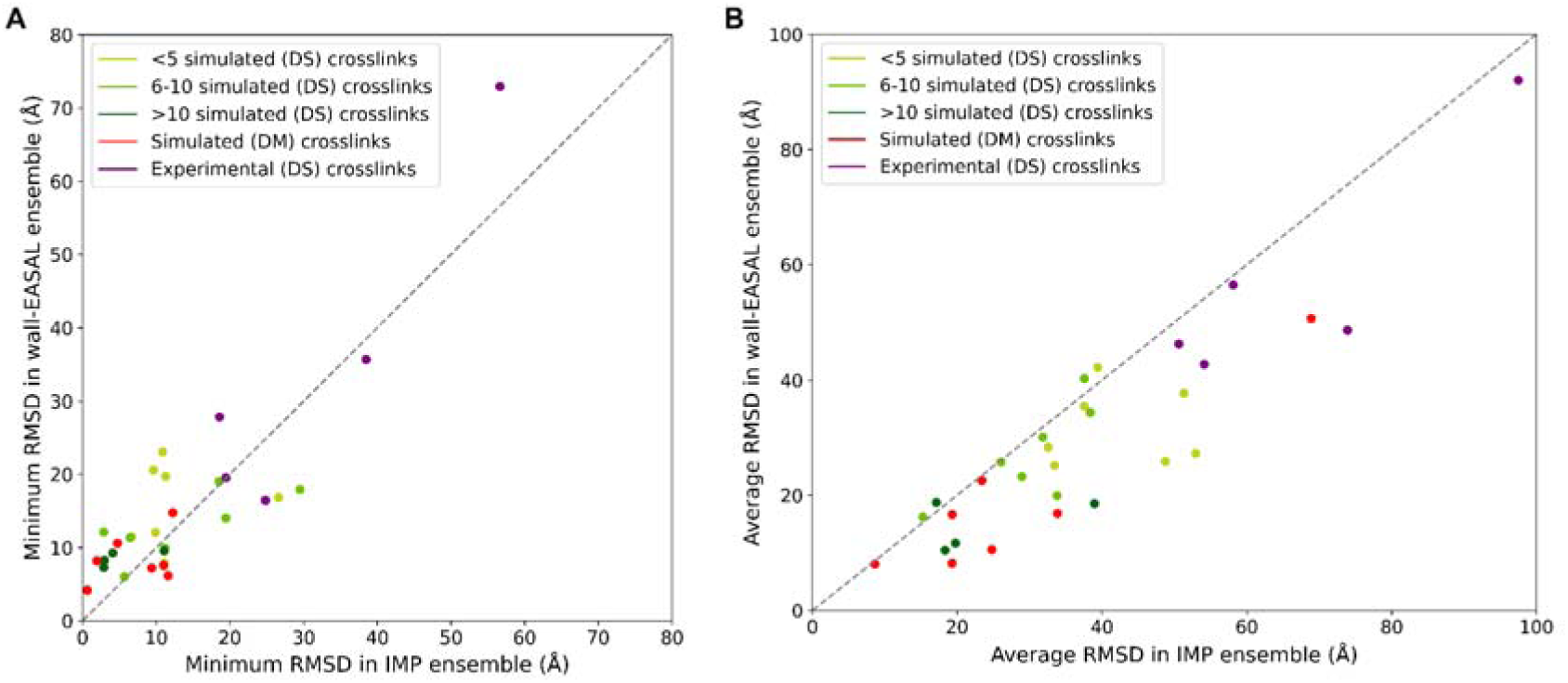
RMSD of the wall-EASAL and IMP configurations. **(A)** Minimum ligand RMSD and **(B)** Average ligand RMSD of a configuration in the IMP and wall-EASAL ensemble to the native structure. DS and DM refer to DSSO and DMTMM crosslinks, respectively.

### Efficiency of IMP and wall-EASAL

IMP relies on randomized sampling that requires multiple independent runs starting from random initial configurations^6,7,17,24^. In contrast, wall-EASAL is a deterministic method that requires a single run per input. The sampling time per independent run for IMP is lower than the time for a wall-EASAL run (Fig. 9A). However, the total sampling time for IMP, in terms of the number of CPU hours, is much higher (Fig. 9B). Moreover, the total runtime for IMP will include the time for analysis, which will add another 25% to the sampling time. Therefore, wall-EASAL is more efficient than IMP.

**Figure 9.**
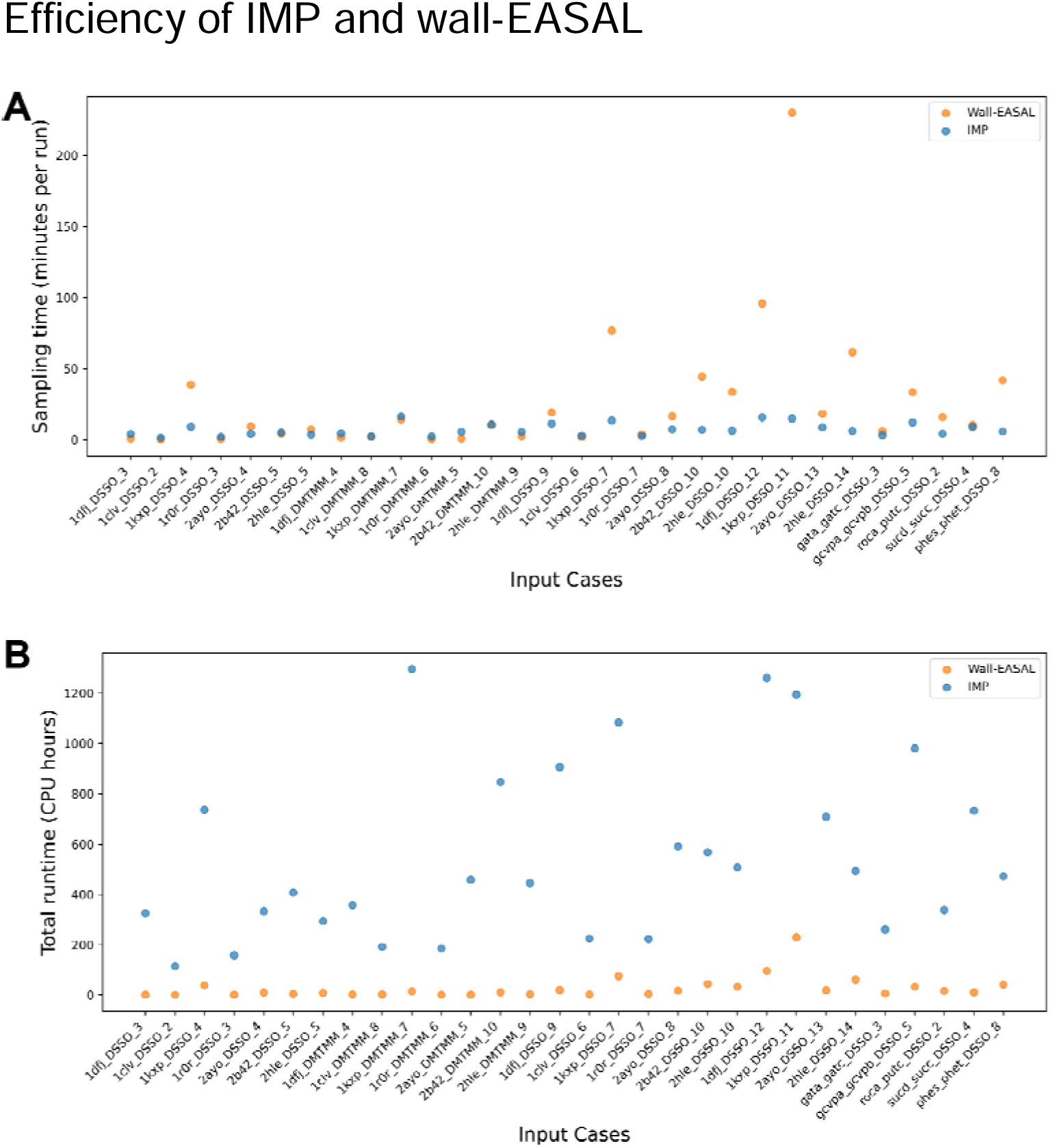
Time efficiency of IMP and wall-EASAL. The distribution of sampling times in terms of the **(A)** average time in CPU minutes per run and **(B)** average total number of CPU hours across the thirty benchmark cases for both methods. All times are on an AMD Ryzen Threadripper 3990x 64-core processor with 256 GB RAM and 2.2 GHz clock speed.

We also compared the efficiency of wall-EASAL and IMP in terms of the number of samples required to obtain the structures that satisfy the input crosslinks sufficiently well. Efficiency was defined by the fraction of structures in the respective ensembles that satisfied the most crosslinks. This comparison was performed for all the cases where the highest percentage of crosslinks satisfied by a single structure in the ensemble was the same for IMP and wall-EASAL. As a general rule, IMP requires many more samples than wall-EASAL to obtain the same maximum crosslink satisfaction as the latter (Fig. 10).

**Figure 10.**
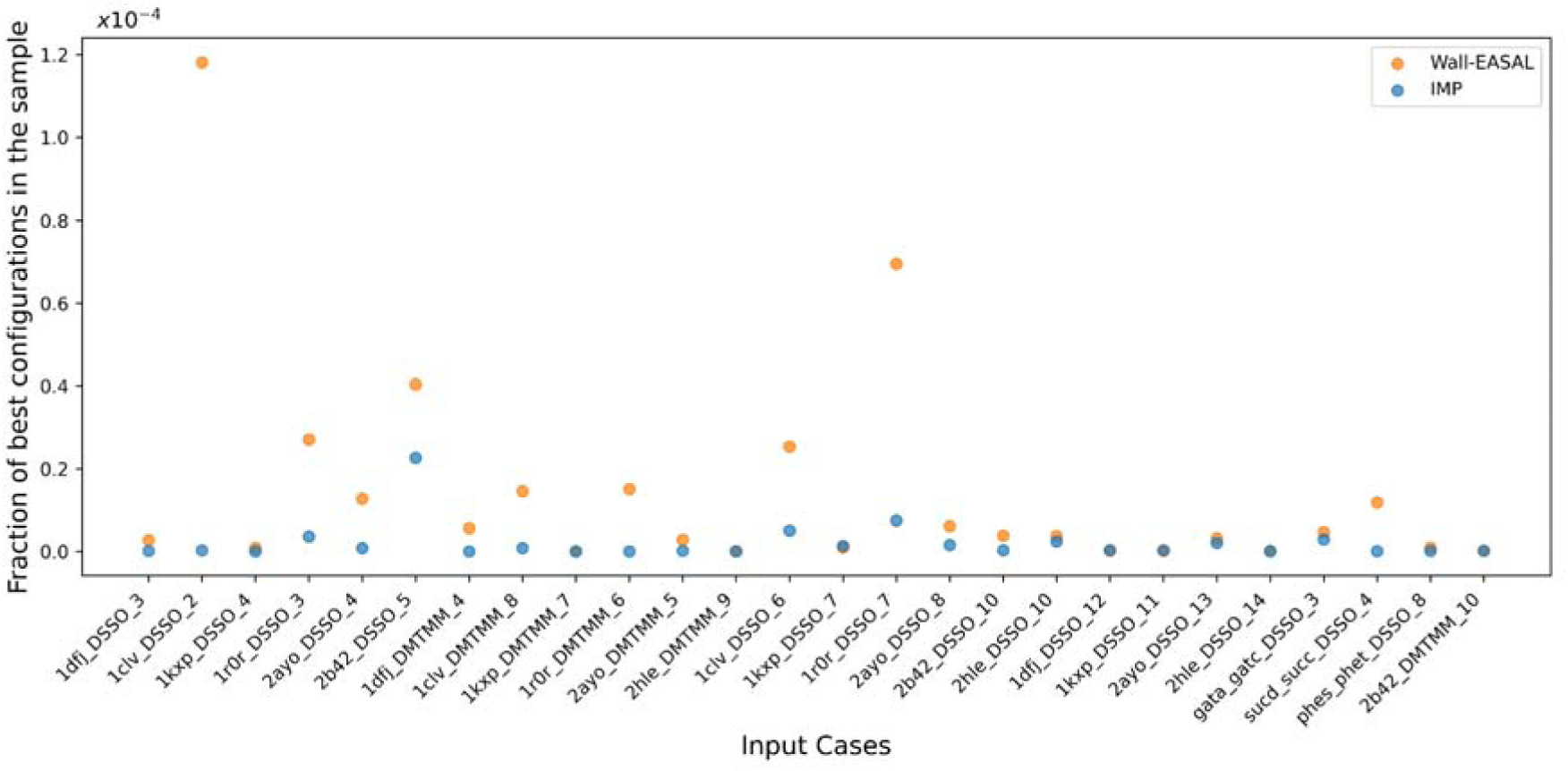
Sampling efficiency of IMP and wall-EASAL. Fraction of configurations in an ensemble with the maximum number of crosslinks satisfied among the total configurations sampled by IMP and EASAL.

For example, in 2hle/DMTMM/9, although the highest percentage of crosslinks satisfied by a configuration is 88% in both ensembles (Fig. 5D, Fig. S2), the fraction of IMP samples that satisfy the maximum number of crosslinks is only 7/100 of the same fraction for wall-EASAL, *i.e.*, the sampling efficiency of wall-EASAL is superior. However, there are exceptions to this rule. For example, in 2b42/DMTMM/10, an IMP configuration satisfies more crosslinks (100%) than a wall-EASAL configuration (90%) (Fig. 5E, Fig. S2), and yet the fraction of wall-EASAL samples that satisfy the maximum number is about the same as for IMP.

### Visualization of structures

Finally, for four input cases, we visualized the best wall-EASAL and IMP configurations, as defined by the configuration with the least ligand RMSD to the corresponding native structure (Fig. 11). For 1dfj/DSSO/3, the best configurations in both the ensembles are similar to the native structure, consistent with our earlier observations that these configurations satisfy the crosslinks well (Fig. 11A). Notably, the crosslink that seems to be long is close to the crosslink upper bound. For 2b42/DMTMM/10, the IMP configuration was closer to the native structure, consistent with the higher crosslink satisfaction observed in the IMP configurations (Fig. 11B, Fig. 5E).

**Figure 11.**
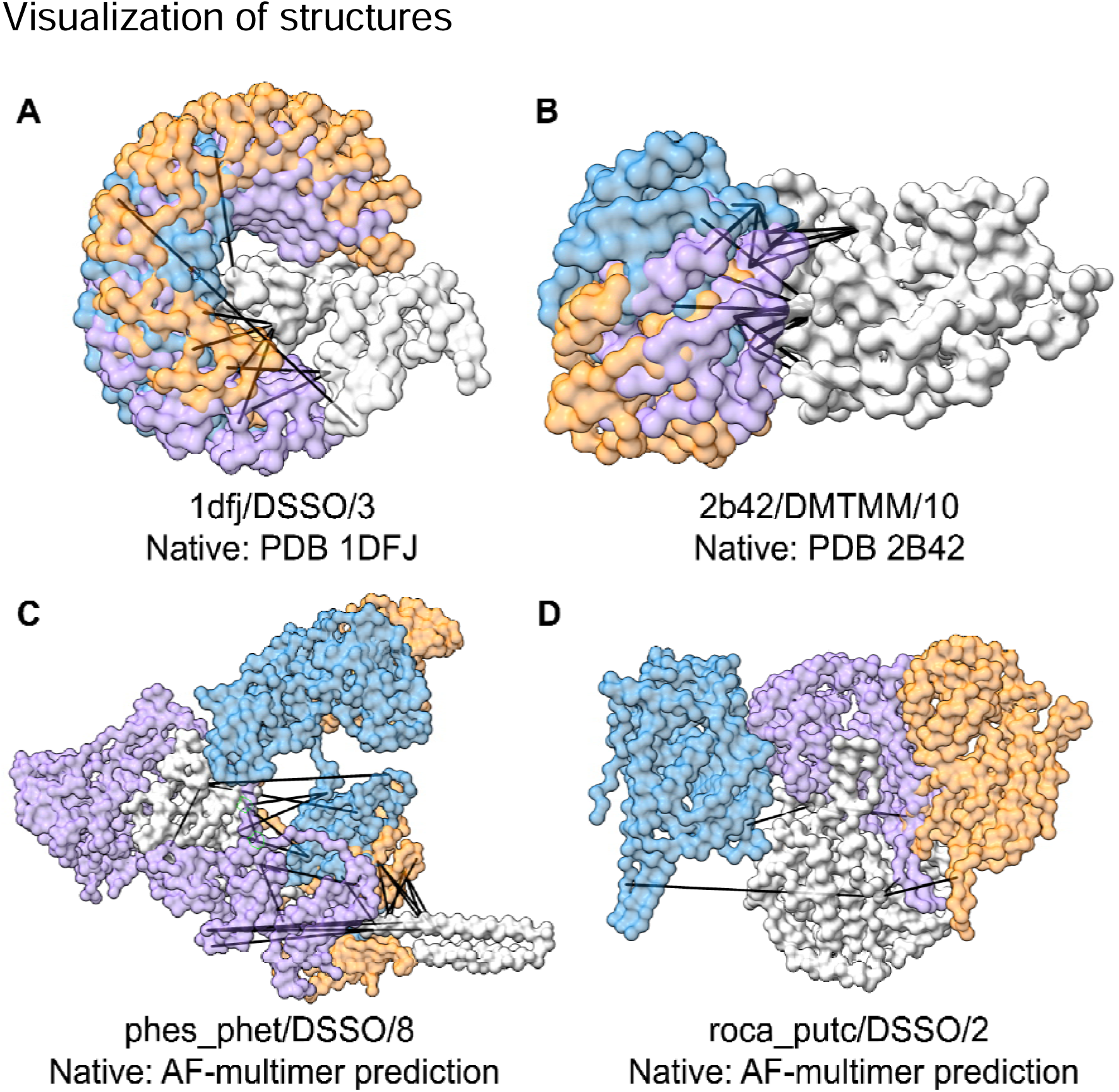
Visualization of wall-EASAL and IMP configurations. The best IMP and wall-EASAL configurations (least ligand RMSD to the native) are superposed on the native structure. **(A-D)** The sampled configuration and native structure are superposed on the receptor (light gray); the ligands in the native structure (purple), wall-EASAL configuration (orange), and IMP configuration (blue) are shown for representative input cases. Crosslinks in the wall-EASAL, IMP, and native configuration are shown by the black lines.

For phes_phet/DSSO/8, both the configurations have large ligand RMSDs from the native structure, *i.e.*, the corresponding AF-multimer prediction. This is intriguing, given that both the ensembles satisfy 87.5% of the crosslinks (Fig. 11C, Fig. S2). This discrepancy might arise because half of the inter-protein crosslinks were violated in the AF-multimer prediction of the phes_phet complex^39^. In this case, our integrative structures are more consistent with the data. For roca_putc/DSSO/2, the wall-EASAL configuration was closer to the native structure, *i.e.*, the corresponding AF-multimer prediction, compared to the IMP configuration, consistent with the earlier observation that the wall-EASAL configurations satisfy the crosslinks better for this case; all crosslinks are also satisfied in the AF-multimer prediction of roca_putc (Fig. 11D, Fig. 5D).

## Discussion

Here, we developed wall-EASAL, a new method for integrative modeling of binary protein-protein complexes given the atomic structures of the constituent proteins and inter-protein chemical crosslinks. The method is based on an efficient discrete geometry algorithm for roadmapping and sampling distance-constrained configurational spaces using distance-based parameterization for dimension reduction and convexification. On a benchmark of thirty input cases, we compared the performance of wall-EASAL with IMP, an integrative modeling method based on randomized sampling. The configurations from wall-EASAL satisfy the crosslinks better as well as resemble the corresponding native structures more closely, on average. Wall-EASAL is also, in general, more efficient than IMP with respect to both measures of efficiency we considered, although there are exceptions to this rule.

Here, we discuss the advantages, uses, limitations, and future directions for integrative docking using wall-EASAL. On the examined benchmark, the method was efficient and produced ensembles that satisfy the input crosslinks well and were close to the native structure. The sampling of configuration space was also demonstrated to be representative, by comparison to the regular EASAL (interior sampling) method for a few input cases. Surprisingly, although at least one crosslink in wall-EASAL’s sample configurations is guaranteed to take an extreme value, in contrast to IMP, wall-EASAL’s distribution of crosslink distances is usually narrower than the IMP distribution of crosslink distances (Fig. 6, Fig. S3).

The assumptions in the method are that the constituent proteins are docked rigidly, their atomic structures are known, and they can be represented by spherical beads coarse-grained at the residue level. Currently, the method as implemented is applicable only for docking of pairs of proteins. In fact, restriction to a protein pair is not a theoretical or algorithmic requirement for the EASAL methodology. The theory behind EASAL encompasses many monomers and many rigid components, some of which could belong to the same monomer and the corresponding algorithmic extension is given in^26^, awaiting implementation. Finally, the crosslink restraint is implemented by a simple distance constraint with upper and lower distance bounds. It does not account for uncertainties in the crosslinking experiment, such as false positive crosslinks^7,44,49^. However, the deterministic distance interval constraint checks can immediately be made probabilistic according to any given noise distribution. Although this could be viewed as more realistic modeling, there is no theoretical guarantee that the deterministic model (more consistent with the Occam’s Razor principle of modeling) would be any less accurate or efficient than the more elaborate probabilistic model.

Here, we discuss parameters that may need to be tuned to improve the performance of wall-EASAL. First, the sampling in wall-EASAL can be made finer (coarser) by reducing (increasing) the step size. We observed that wall-EASAL can get solutions with a step size of 20 (‘*stepSize*’), however, for a few input cases using a lower step size of 5 was required to find the best configurations. There is a trade-off between the step size and the sampling time. The sampling time could increase up to 2.5 fold upon reducing the step size by half. Second, one may need to decrease the crosslink satisfaction tolerance (*‘crossLinkSatisfyThres*’) value if configurations that satisfy the specified number of crosslinks are not found.

It is conceivable to introduce walls at intermediate distances (*‘smartCrosslinkMode*’) in the interior of the crosslink interval to provide a larger configurational space for wall-EASAL to sample configurations from. However, additional walls in the interior would result in a corresponding decrease in efficiency with questionable returns since we have demonstrated that in the absence of pockets or links or their coarse sampling artifacts, wall-EASAL guarantees a feasible solution satisfying all (or the maximum possible number) of the crosslinks, and moreover provides a representative collection of configurations of the entire feasible region including the interior.

Any information on distances between the residues or domains of constituent proteins can be used in wall-EASAL; although the current study uses chemical crosslinks, other types of distances can also be used, for example from NMR or genetic interaction assays^50^. Structures of constituent proteins can be derived from experiments or AI-based predictions^51^. The method may be of particular interest in cases where the structures of constituent proteins have not been experimentally determined, but reliable AlphaFold predictions of the monomer are available, along with crosslinks^39,52,53^. Structures of antibody-antigen complexes are also of special interest since AI-based predictions of these complexes are not currently reliable^54–56^.

The structures of binary complexes from wall-EASAL can complement methods for integrative modeling of macromolecular assemblies. For instance, these structures can suggest rigid bodies or restraints on protein interfaces for use in IMP, Assembline, or Haddock^6,14–17^. Such information on pairs of proteins can then be combined with other information to model a larger complex. The structures from wall-EASAL can also be used as inputs for methods that perform combinatorial searches for structures of large assemblies based on the component binary complexes, such as CombFold and MCTreeSearch^32–34^.

New geometric deep learning methods that predict the optimal distance between crosslinked residues can be used to further refine the inputs to methods such as wall-EASAL^57^. Future planned extensions of the method include parallelizing it for efficiency and modifying the algorithm to scale to larger constituent proteins and protein complexes with multiple subunits. Bayesian formulations of restraints can be used instead of simple distance restraints^44^. The method can be extended to include restraints other than pairwise distance restraints, such as EM-based shape restraints, by devising ways to convert them to equivalent distance restraints when possible. Incorporating wall-EASAL in integrative modeling methods such as IMP will facilitate the characterization of assemblies and cellular neighborhoods at increased efficiency, accuracy, and precision.

## Data and software availability

The implementation of the wall-EASAL method is available at https://bitbucket.org/geoplexity/easal-dev/src/Crosslink. The integrative docking benchmark is available at https://github.com/isblab/Integrative_docking_benchmark. The benchmark is also available at Zenodo: https://doi.org/10.5281/zenodo.13959115.

## Supporting information

Supporting information contains figures showing the input structures (Fig. S1), percentage of crosslink satisfaction (Fig. S2), average crosslink distance (Fig. S3), crosslink distance difference in the sampled configurations and the native structure (Fig. S4), and RMSD of the wall-EASAL and IMP configurations (Fig. S5) in each benchmark case. Table S1 contains the description of the benchmark. The Mathematical proof that wall-EASAL finds a feasible configuration satisfying crosslink constraints if one exists is also given.

## Acknowledgments

We thank ISB Lab members Shreyas Arvindekar, Kartik Majila, Omkar Golatkar, and Mubashira KP for their useful comments on the manuscript. Molecular graphics images were produced using the UCSF Chimera and UCSF ChimeraX packages from the Resource for Biocomputing, Visualization, and Informatics at the University of California, San Francisco (supported by NIH P41 RR001081, NIH R01-GM129325, and National Institute of Allergy and Infectious Diseases). The authors acknowledge University of Florida’s UFIT Research Computing for providing the Hipergator computational resources that have contributed to the research results reported in this publication.

---

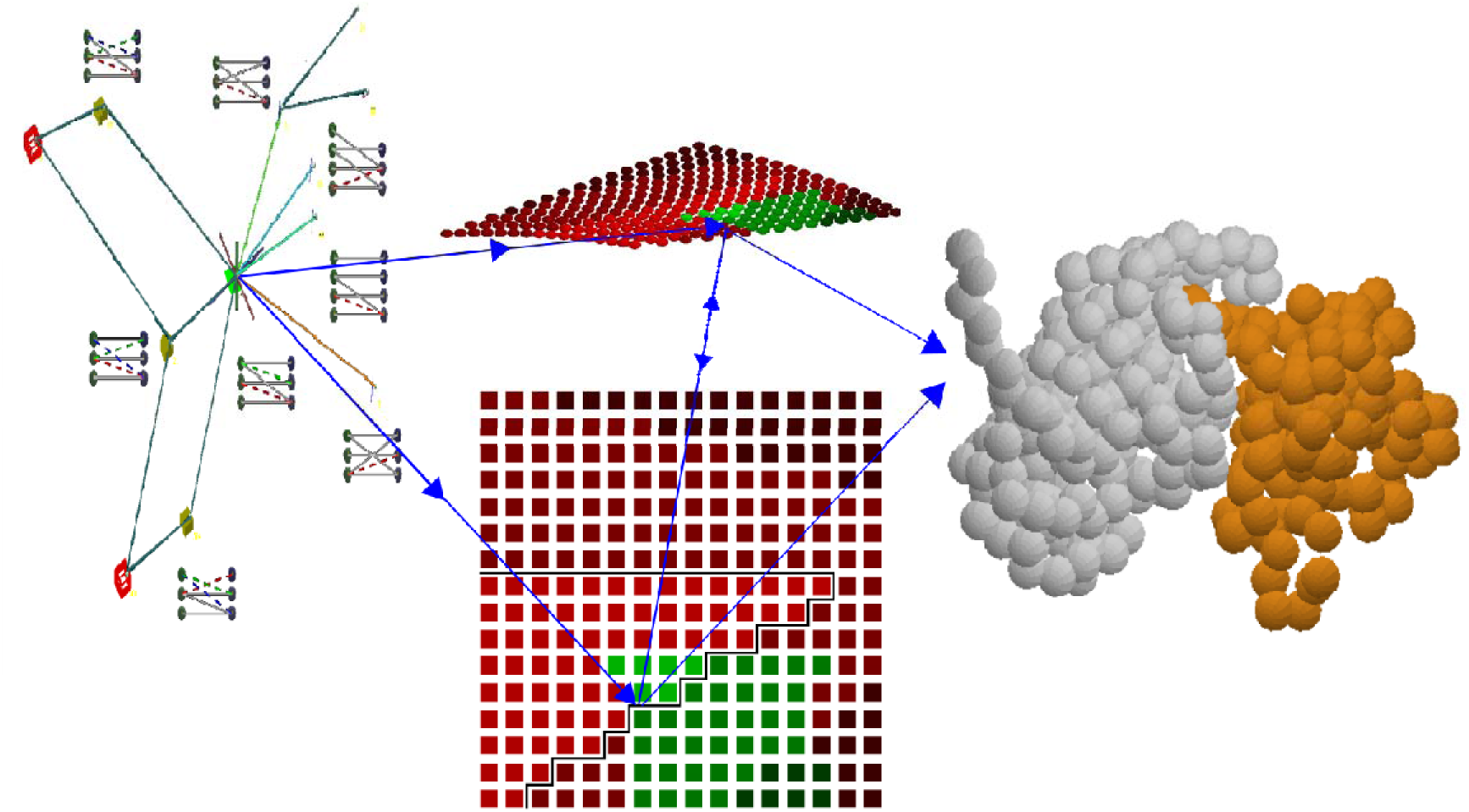
Legend: Given input structures of two proteins and chemical crosslinks between them, Wall-EASAL optimizes sampling to guarantee an output set of representative configurations of the complex satisfying the input constraints.

