## Supplementary Material for "A new discrete-geometry approach for integrative docking of proteins using chemical crosslinks"

### Supplementary Figures

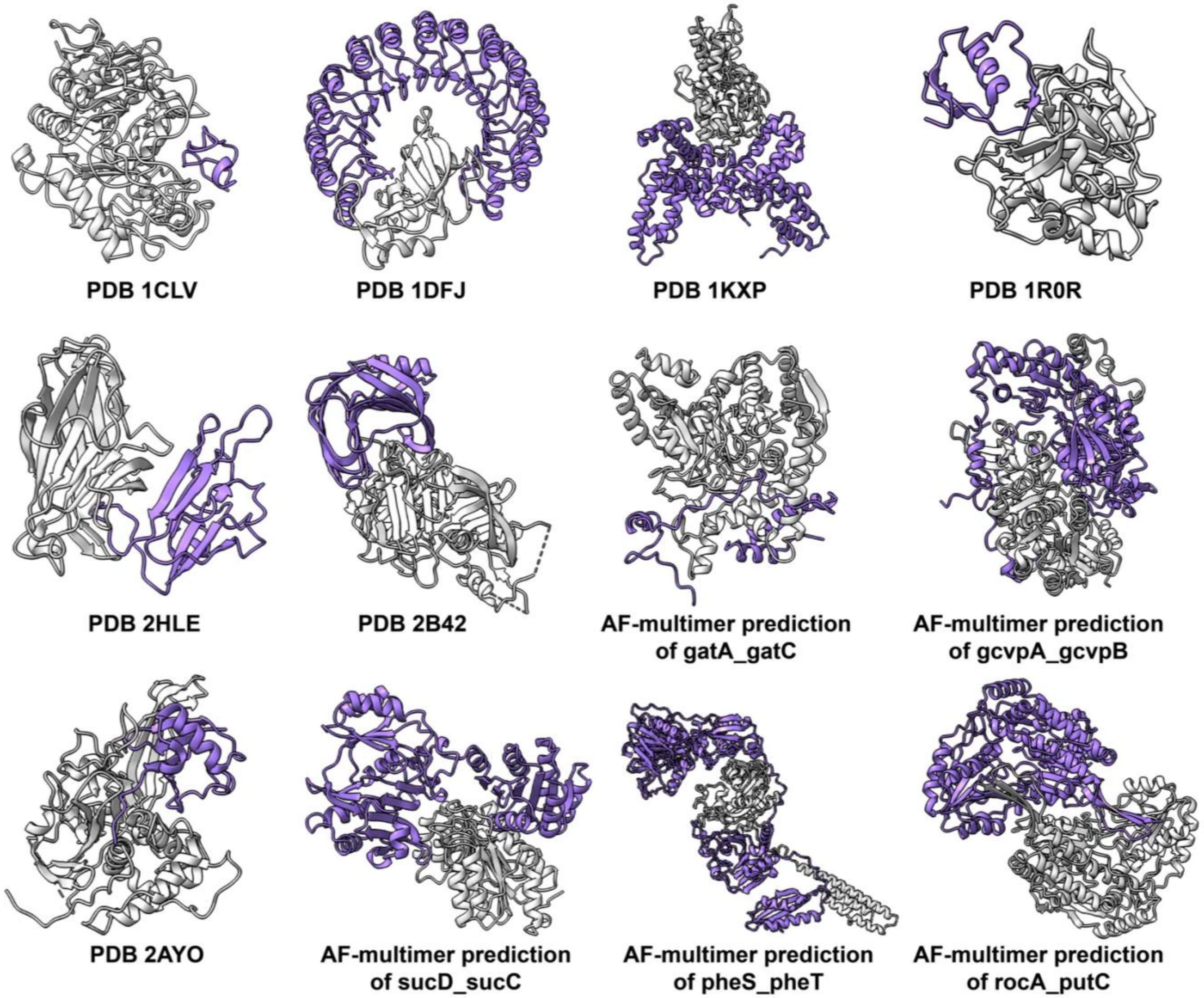

**Figure S1. Structures of the binary complexes.** Seven PDBs and five AlphaFold-multimer predicted complexes were used as benchmark inputs for integrative docking, obtained from Zlab benchmark 5.5 (Guest et al., 2021) and O'Reilly, Molecular Systems Biology, 2023 (O'Reilly et al., 2023). The receptor and ligand are shown in light grey and purple, respectively.

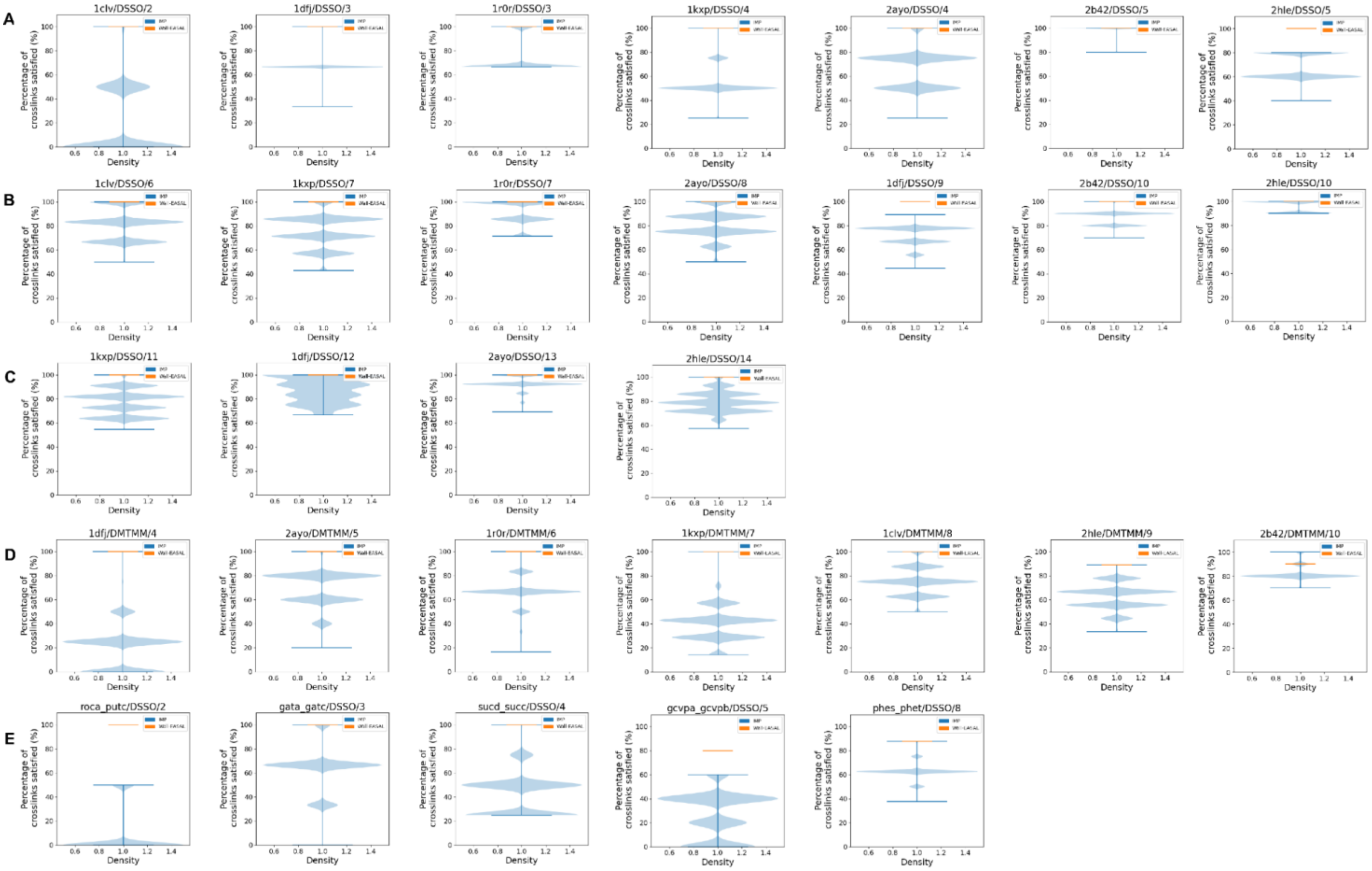

**Figure S2. Percentage of crosslinks satisfied in wall-EASAL and IMP ensembles. (A)** Input cases with five or fewer, **(B)** between six and ten, and **(C)** ten or more simulated DSSO crosslinks. **(D)** Input cases with simulated DMTMM crosslinks. **(E)** Input cases with DSSO crosslinks from experiments. The monomer structures were derived from the structure of the complex in the PDB **(A-D)** or were predicted by Alphafold2 **(E)** (Table S1) (Jumper et al., 2021).

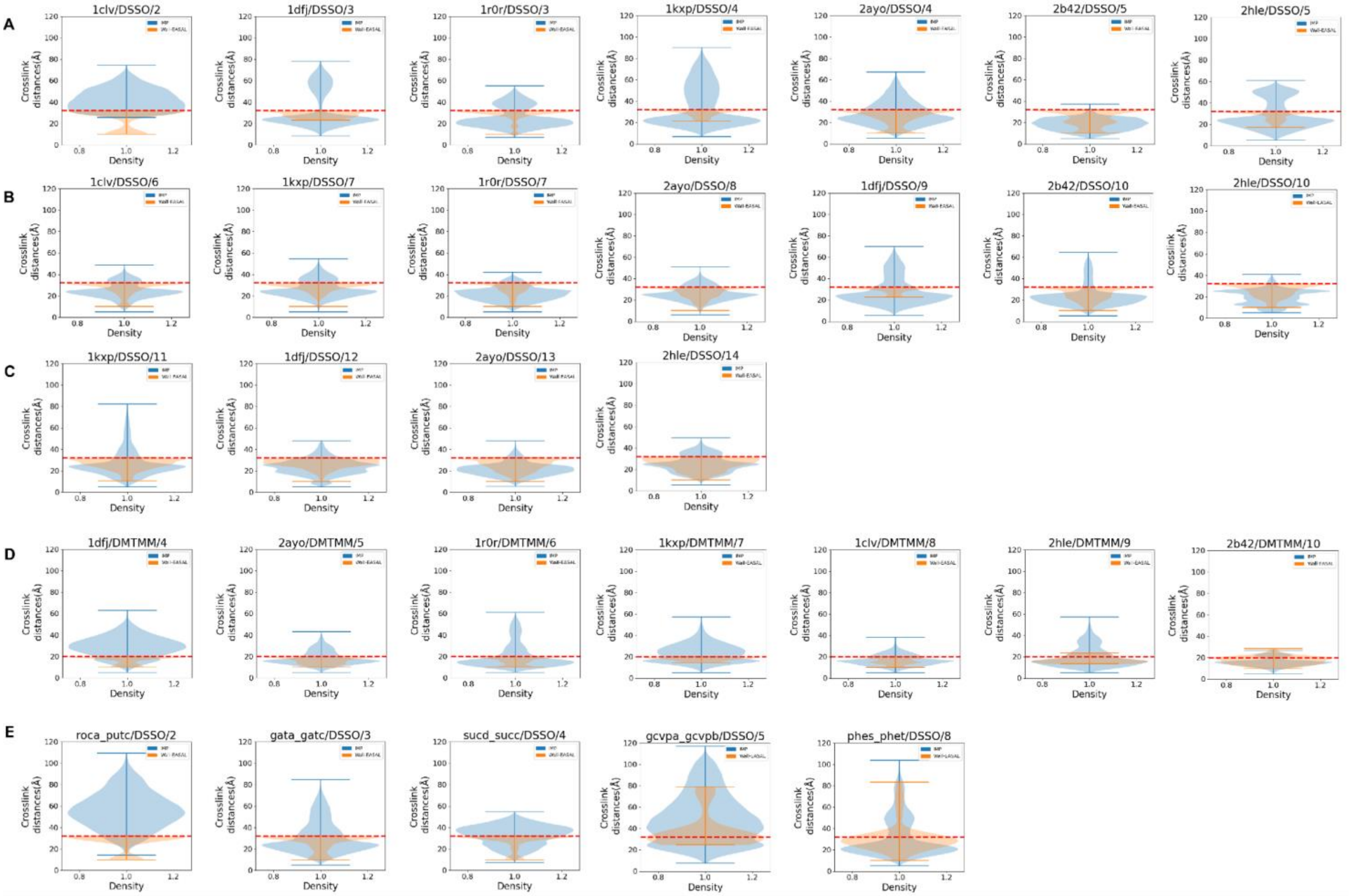

**Figure S3. Distribution of crosslink distance in wall-EASAL and IMP ensembles. (A)** Input cases with five or fewer, **(B)** between six and ten, and **(C)** ten or more simulated DSSO crosslinks. **(D)** Input cases with simulated DMTMM crosslinks. **(E)** Input cases with DSSO crosslinks from experiments. The monomer structures were derived from the structure of the complex in the PDB **(A-D)** or were predicted by Alphafold2 **(E)** (Table S1) (Jumper et al., 2021).

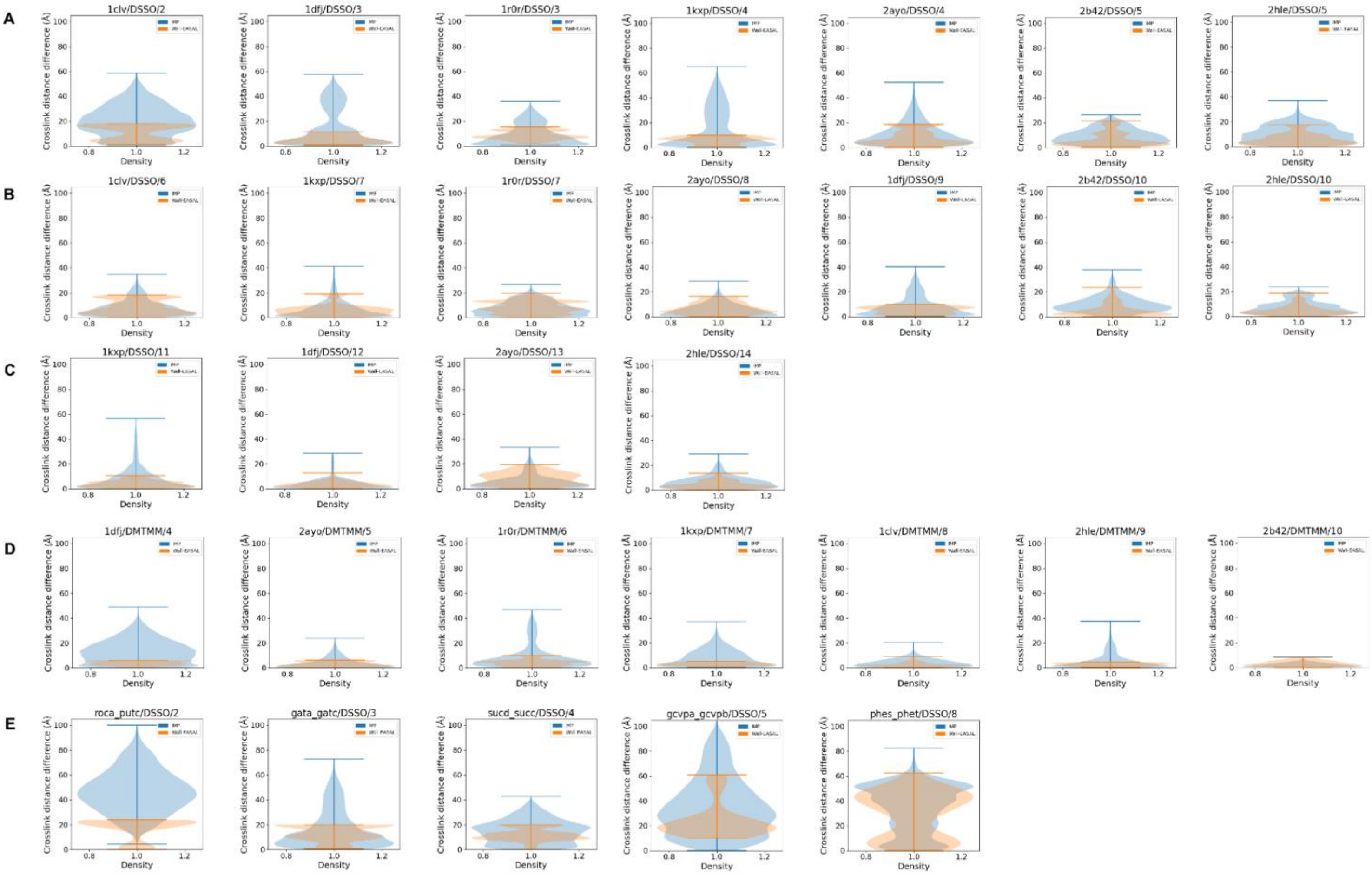

**Figure S4. Comparison between crosslink distances in the sampled configurations and the native structure. (A)** Input cases with five or fewer, **(B)** between six and ten, and **(C)** ten or more simulated DSSO crosslinks. **(D)** Input cases with simulated DMTMM crosslinks. **(E)** Input cases with DSSO crosslinks from experiments. The monomer structures were derived from the structure of the complex in the PDB **(A-D)** or were predicted by Alphafold2 **(E)** (Table S1) (Jumper et al., 2021).

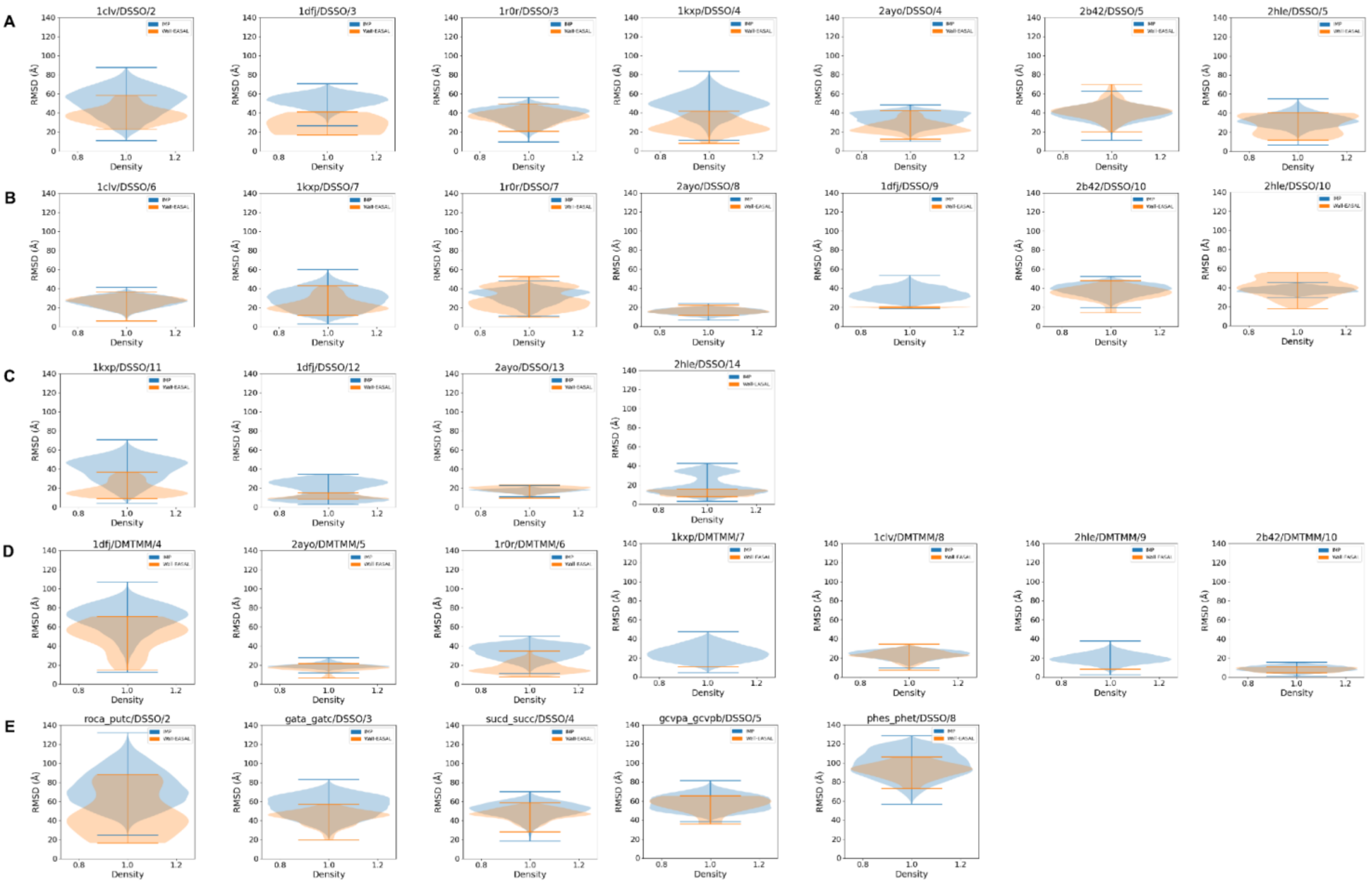

**Figure S5. RMSD of wall-EASAL and IMP sampled configurations to the native structure.** **(A)** Input cases with five or fewer, **(B)** between six and ten, and **(C)** ten or more simulated DSSO crosslinks. **(D)** Input cases with simulated DMTMM crosslinks. **(E)** Input cases with DSSO crosslinks from experiments. The monomer structures were derived from the structure of the complex in the PDB **(A-D)** or were predicted by Alphafold2 **(E)** (Table S1) (Jumper et al., 2021).

### Supplementary Table

| Categories based on the source of crosslinks | Number of crosslinks | Complex name or PDB ID | Receptor, ligand chain | Source of monomer structure (Experiment *i.e.* PDB or AlphaFold-predicted) | Reference | Type |
|---|---|---|---|---|---|---|
| <5 Simulated DSSO crosslinks | 2 | 1clv | A,I | PDB | Zlab benchmark 5.5 | EI |
| | 3 | 1dfj | E,I | PDB | Zlab benchmark 5.5 | EI |
| | 4 | 1kxp | A,D | PDB | Zlab benchmark 5.5 | OX |
| | 3 | 1r0r | E,I | PDB | Zlab benchmark 5.5 | EI |
| | 4 | 2ayo | A,B | PDB | Zlab benchmark 5.5 | ER |
| | 5 | 2b42 | A,B | PDB | Zlab benchmark 5.5 | EI |
| | 5 | 2hle | A,B | PDB | Zlab benchmark 5.5 | OR |
| 6-10 Simulated DSSO crosslinks | 6 | 1clv | A,I | PDB | Zlab benchmark 5.5 | EI |
| | 9 | 1dfj | E,I | PDB | Zlab benchmark 5.5 | EI |
| | 7 | 1kxp | A,D | PDB | Zlab benchmark 5.5 | OX |
| | 7 | 1r0r | E,I | PDB | Zlab benchmark 5.5 | EI |
| | 8 | 2ayo | A,B | PDB | Zlab benchmark 5.5 | ER |
| | 10 | 2b42 | A,B | PDB | Zlab benchmark 5.5 | EI |
| | 10 | 2hle | A,B | PDB | Zlab benchmark 5.5 | OR |
| >10 Simulated DSSO crosslinks | 12 | 1dfj | E,I | PDB | Zlab benchmark 5.5 | EI |
| | 11 | 1kxp | A,D | PDB | Zlab benchmark 5.5 | OX |
| | 13 | 2ayo | A,B | PDB | Zlab benchmark 5.5 | ER |
| | 14 | 2hle | A,B | PDB | Zlab benchmark 5.5 | OR |
| Simulated DMTMM crosslinks | 8 | 1clv | A,I | PDB | Zlab benchmark 5.5 | EI |
| | 4 | 1dfj | E,I | PDB | Zlab benchmark 5.5 | EI |
| | 7 | 1kxp | A,D | PDB | Zlab benchmark 5.5 | OX |
| | 6 | 1r0r | E,I | PDB | Zlab benchmark 5.5 | EI |
| | 5 | 2ayo | A,B | PDB | Zlab benchmark 5.5 | ER |
| | 10 | 2b42 | A,B | PDB | Zlab benchmark 5.5 | EI |

| | | | | | | |
|---|---|---|---|---|---|---|
| | 9 | 2hle | A,B | PDB | Zlab benchmark 5.5 | OR |
| DSSO crosslinks from experiments | 3 | gata-gatc | A,B | AF-multimer | O'Reilly *et al*, Molecular Systems Biology, 2023 | ER |
| | 5 | gcvpa-gcvpb | A,B | AF-multimer | O'Reilly *et al*, Molecular Systems Biology, 2023 | ER |
| | 8 | phes-phet | A,B | AF-multimer | O'Reilly *et al*, Molecular Systems Biology, 2023 | ER |
| | 2 | roca-putc | A,B | AF-multimer | O'Reilly *et al*, Molecular Systems Biology, 2023 | ER |
| | 4 | sucd-succ | A,B | AF-multimer | O'Reilly *et al*, Molecular Systems Biology, 2023 | ER |

**Table S1: Benchmark dataset.** The dataset is categorized based on the source and the number of crosslinks. There are five categories: five or fewer, between six and ten, and ten or more simulated DSSO crosslinks, simulated DMTMM crosslinks, and DSSO crosslinks from experiments. There are also five categories of the complexes based on the type of protein: enzyme–inhibitor (EI); enzyme–substrate (ES); enzyme complex with a regulatory or accessory chain (ER); others, receptor containing (OR); others, miscellaneous (OX) (Guest et al., 2021). The monomer structures are obtained from Zlab benchmark 5.5 (Guest et al., 2021) and O'Reilly, Molecular Systems Biology, 2023 (O’Reilly et al., 2023).

### Mathematical proof that wall-EASAL finds a feasible configuration satisfying crosslink constraints if one exists

#### Problem Description

Given:

- Two point-sets $A = \{A_1, A_2, ..., A_m\}$ , $B = \{B_1, B_2, ..., B_n\}$,
- A non-empty Constraint Graph $\boldsymbol{G} = (\boldsymbol{V} \subseteq \boldsymbol{A} \cup \boldsymbol{B}, \boldsymbol{E})$: The edge $\boldsymbol{e} \in \boldsymbol{E}$ represents a crosslink, which is a distance (interval) constraint between endpoints of $\boldsymbol{e} = (\boldsymbol{v}, \boldsymbol{w})$ where $\boldsymbol{v} \in \boldsymbol{A}, \boldsymbol{w} \in \boldsymbol{B}$.
- Variables of the system are Euclidean isometries $T_A, T_B$, whose instantiations are the *configurations*
- The distance interval **constraints**:
  - **C1(collision)**: $\forall v \in A, w \in B, l(v,w) \leq ||T_A(v) - T_B(w)||, l \in R^+$
  - **C2(crosslink)**:$\forall (v,w) \in E(G), l(v,w) \leq ||T_A(v) - T_B(w)|| \leq h(v,w), h \in R^+$
  - **C3(wall)**:$\exists (v,w) \in E(G), ||T_A(v) - T_B(w)|| = l(v,w)\ or\ ||T_A(v) - T_B(w)|| = h(v,w)$

The goal is to obtain a solution satisfying all constraints, showing that the addition of **C3** to the system does not affect the existence of solutions.

*Theorem*

Let $R_S{}^1$ be the set of configurations satisfying **C1**. Let $R_S$ be the configuration space satisfying **C1** and **C2**, and $R_S'$, the *wall* configuration space satisfying **C1**, **C2**, and **C3**,
If $R_S{}^1$ is path-connected. then $R_S$ is non-empty if and only if $R_S'$ is non-empty.

*Proof:* Let $R_S{}^2$ be the set of configurations satisfying **C2** and $R_S{}^{2*}$ be the set of configurations satisfying **C2** and **C3**. Notice that $R_S{}^1$, $R_S{}^2$, $R_S{}^{2*}$ are all closed sets. Since arbitrarily large transformations $T$ satisfy **C1**, $R_S{}^1$ is unbounded. $G$ has at least one edge, and $h(v,w)$ is finite, so $R_S{}^2$ is bounded. Furthermore, $R_S{}^{2*}$ is exactly the boundary of $R_S{}^2$, denoted $\Omega(R_S{}^2)$. Therefore $R_S = R_S{}^1 \cap R_S{}^2$ and $R_S' = R_S{}^1 \cap R_S{}^{2}{}'$. A simple case is when $R_S{}^2$ is not full dimensional, then it has an empty interior, thus $R_S{}^2 = R_S{}^{2}{}'$ and $R_S = R_S'$, proving the theorem.

In general, the non-empty intersection of any closed and bounded set $U$ with a closed, connected set $W$ contains the nonempty intersection of $W$ with $\Omega(U)$ unless $W \subsetneq U$, which is impossible if $W$ is unbounded. Now the theorem is proven by substituting $U = R_S{}^2, \Omega(U) = \Omega(R_S{}^2), W = R_S{}^1$.
